# mbkmeans: fast clustering for single cell data using mini-batch *k*-means

**DOI:** 10.1101/2020.05.27.119438

**Authors:** Stephanie C. Hicks, Ruoxi Liu, Yuwei Ni, Elizabeth Purdom, Davide Risso

**Affiliations:** Department of Biostatistics, Johns Hopkins Bloomberg School of Public Health, MD, USA; Department of Applied Mathematics and Statistics, Johns Hopkins University, Baltimore, MD, USA; Division of Biostatistics and Epidemiology, Department of Healthcare Policy and Research, Weill Cornell Medical College, New York, NY, USA; Department of Statistics, University of California, Berkeley, Berkeley, CA, USA; Department of Statistical Sciences, University of Padua, Italy

## Abstract

Single-cell RNA-Sequencing (scRNA-seq) is the most widely used high-throughput technology to measure genome-wide gene expression at the single-cell level. One of the most common analyses of scRNA-seq data detects distinct subpopulations of cells through the use of unsupervised clustering algorithms. However, recent advances in scRNA-seq technologies result in current datasets ranging from thousands to millions of cells. Popular clustering algorithms, such as *k*-means, typically require the data to be loaded entirely into memory and therefore can be slow or impossible to run with large datasets. To address this problem, we developed the *mbkmeans* R/Bioconductor package, an open-source implementation of the mini-batch *k*-means algorithm. Our package allows for on-disk data representations, such as the common HDF5 file format widely used for single-cell data, that do not require all the data to be loaded into memory at one time. We demonstrate the performance of the *mbkmeans* package using large datasets, including one with 1.3 million cells. We also highlight and compare the computing performance of *mbkmeans* against the standard implementation of *k* - means. Our software package is available in Bioconductor at https://bioconductor.org/packages/mbkmeans.

## Introduction

Unsupervised clustering algorithms are commonly used in areas such as genomics, machine learning, pattern recognition, and data compression to divide a set of unlabeled observations into separate groups with similar traits [1, 2]. In particular, clustering algorithms are popular in single-cell transcriptomics, where datasets can consist of millions of unlabeled observations (or cells) [3, 4]. The goal in this setting is to group cells into distinct clusters with discrete labels that approximate true biological groups [5]. In this context, different clusters can be thought of as different cell types or cell states, which can be further explored in downstream analyses [4].

The most widely used partitional clustering algorithm is *k*-means [6, 7, 8]. The algorithm partitions *N* cells into *k* clusters each represented by a centroid, or mean profile, for the cells in the *k*^*th*^ cluster. This algorithm is commonly used not only on its own, but also as a component of ensemble clustering [9, 10].

While *k*-means is easy to implement and can be fast for small enough data, it assumes that the user has enough computational resources (specifically RAM) to store the data (and Euclidean distances between observations and centroids) into memory. However, file sizes generated from scRNA-seq experiments can be on the order of tens to hundreds of gigabytes. For large enough data, *k*-means can be slow or completely fail if a user lacks sufficient computational resources. Ensemble clustering approaches that depend on the use of *k*-means [9, 10] run it multiple times (e.g., with different parameter values or on a different data subset) limiting the usability of these packages for large scRNA-seq datasets [11]. We note that our goal here is not to debate the relative merits of *k*-means as a clustering algorithm – *k*-means is a well-established method, which has been thoroughly investigated [12] – but to provide users with the ability to use the popular *k*-means algorithm on large single-cell datasets.

To address the problems of using *k*-means with large data, two solutions are (1) parallelization and (2) subsampling. Parallelization approaches typically leverage some combination of (i) *MapReduce* [13] *concepts to handle a large volume of data over a distributed computing environment [14, 15], (ii) k*-dimensional (*k*-d) trees to either optimize for the nearest centroid [16] or to partition datasets into subsets, representative of the larger dataset [17], and (iii) leverage multi-core processors [18]. While these approaches do improve the speed of *k*-means, they can be limited to the number of reducers for each centroid and can often require extensive computational resources. In contrast, subsampling approaches, such as the mini-batch *k*-means algorithm [19] work on small, random subsamples of data (“mini batches”) that can fit into memory on standard computers.

Current implementations of the mini-batch *k*-means algorithm [19] are available in standard programming languages such as in the *scikit-learn* machine learning Python library [20] or in the *ClusterR* R package [21]. However, these implementations either implicitly or explicitly require all the data to be read into memory, and therefore do not leverage the potential of the algorithm to provide a low memory footprint.

To address the described problems, we implemented the <u>m</u>ini-<u>b</u>atch *<u>k</u>*<u>-means</u> clustering algorithm in the open-source *mbkmeans* R package [22], providing fast, scalable, and memory-efficient clustering of scRNA-seq data in the Bioconductor framework [5, 23]. Like existing implementations, our package can be applied to in-memory data input (useful for smaller datasets). However, our implementation can also work with on-disk data, taking advantage of the subsampling structure of the algorithm to read into memory only the current “mini batch” of data at any given point, thereby greatly reducing the required memory (RAM) needed. *mbkmeans* accomplishes this task by directly accessing the needed data for any mini-batch from data stored in an on-disk format, such as the HDF5 file format [24], which is widely used for distributing single-cell sequencing data.

We evaluate the performance of *mbkmeans* (both on in-memory and on-disk data) compared to the standard *k*-means (in-memory data only) algorithm using the 1.3 million brain cells scRNA-seq data from 10X Genomics and using simulation studies. We demonstrate that our implementation constrains the memory usage of the clustering algorithm, increasing speed and without loss of accuracy as compared to the standard *k*-means algorithm. Our contribution is two-fold: we implement a mini-batch *k*-means algorithm for on-disk data, and we benchmark the performance of a non-trivial algorithm for HDF5 against its in-memory counterpart.

## Design and Implementation

Broadly, the mini-batch *k*-means algorithm works in a similar spirit to standard *k*-means. The primary difference is that at each iteration, the algorithm uses a small, random subset (or a mini-batch) of a fixed size *b*, so that the subset can be loaded in memory. First, we briefly describe the *k*-means algorithm, and then we contrast it with the mini-batch *k*-means algorithm [19]. Throughout, we use `teletype` font when referring to specific code or functions, and *italicized* font when referring to software packages.

### Overview of *k*-means algorithm

Given a set of observations **Y** = (**y**_1_, **y**_2_, …, **y**_*N*_) where each observation is a *G*-dimensional real vector, the optimization problem of *k*-means clustering is to partition the *N* observations into *k*(≤ *N*) sets **S** = {*S*_1_, *S*_2_, …, *S*_*k*_} to minimize the within-cluster sum of squares (WCSS) or

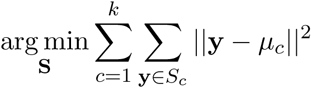

where *µ*_*c*_ is the centroid of observations in *S*_*c*_. Lloyd’s algorithm [8] is the most widely used algorithm to solve this optimization, alternating between an **assignment step** and an **update step** until convergence.

### Overview of mini-batch *k*-means algorithm

While *k*-means is widely used, two problems with its implementation are (1) it requires all the data to be stored in memory and (2) it may be computationally expensive when the sample size and/or the number of clusters are large. Stochastic gradient descent methods compute a gradient descent step one observation at a time, and are often used to improve the speed and memory usage [26]. The primary disadvantage is they find typically lower quality solutions due to stochastic noise; a standard solution is to use batches of data at each gradient descent, rather than individual data points (mini-batch gradient descent). This strategy was proposed in the context of *k*-means [19] for improving its computational performance, and is known as the mini-batch *k*-means algorithm. The use of mini-batches has been shown to have lower stochastic noise without the expensive computational time in the *k*-means algorithm with large datasets [19]. The use of the mini-batches also allows for the ability to not store the entire dataset in memory. Therefore, the mini-batch *k*-means algorithm is an ideal approach to allow for fast, scalable, memory-efficient clustering of scRNA-seq data.

The *mbkmeans* follows a similar iterative approach to Lloyd’s algorithm. However, at each iteration *t*, a new random subset of size *b*, specifically **M**, is used and this continues until convergence. If we define the number of centroids as *k* and the mini-batch size as *b* (what we refer to as the ‘batch size’), then our implementation of mini-batch *k*-means follows that of *ClusterR* [21], *and is briefly described here:*

0. *At t* = 0: Initialize the set of *k* centroids 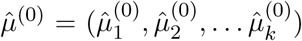.
1. For each *t* ≥ 1: Randomly sample (without replacement) from **Y** a random subset **M** of size *b*. Update the estimates of the *k* centroids by performing the following two steps:
  i. **Assignment**: Given the set of *k* centroids at *t* − 1, or 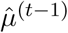, compute the Euclidean distances between observations in **M** and the *k* cluster centroids. Assign each observation from **M** to the closest centroid to obtain a new set of observations per centroid 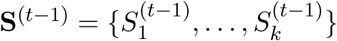.
  ii. **Update**: calculate the new centroids by averaging the coordinates of the observations from the mini-batch **M** assigned to each cluster to obtain 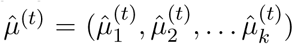.
  Steps (i-ii) are repeated until convergence using the difference in norm of the centroids, 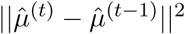.
2. While Step 1 returns a set of estimated *k* cluster centroids, most often we are interested in assigning a label to each observation. This is automatically returned in *k*-means clustering with Lloyd’s algorithm, which uses all the observations to estimate the centroids. However, mini-batch *k*-means uses the final estimates of the *k* centroids in a final prediction step to obtain cluster labels of all observations in **Y**.

### The *mbkmeans* software package

The *mbkmeans* software package implements the mini-batch *k*-means clustering algorithm described above and works with matrix-like objects as input. Specifically, the package works with standard R data formats that store the data in memory, such as the standard *matrix* class in base R and sparse and dense matrix classes from the *Matrix* R package [27], and with file-backed matrices, e.g., by using the HDF5 file format [24]. In addition, the package provides methods to interface with standard Bioconductor data containers such as the *SummarizedExperiment* [28] *and SingleCellExperiment* [29] classes.

We implemented the computationally most intensive steps of our algorithm in C`++`, leveraging the *Rcpp* and *beachmat* [31] packages. Furthermore, we make use of Bioconductor’s *DelayedArray* [32] framework, and in particular the *HDF5Array* [33] package to interface with HDF5 files. The *mbkmeans* package was built in a modular format that would allow it to easily operate on alternative on-disk data representations in the future. To initialize the *k* centroids, the *mbkmeans* package uses the *k-means*++ initialization algorithm [34] with a random subset of *b* observations (the batch size), by default. Finally, to predict final cluster labels, we use block processing through the *DelayedArray* [32] package to avoid working with all the data at once, which could be problematic if the dataset does not entirely fit in memory.

### Benchmarking datasets

To evaluate the performance of *mbkmeans*, we used: (i) simulated gene expression data from a mixture of *k* Gaussian distributions, and (ii) downsampled subsets of a real scRNA-seq dataset from 10X Genomics [25].

We evaluate both the memory usage and computing time using the `Rp`r`of` and `proc.time` R functions. Both of this are reported from two independent computing systems: (i) an iMac with a 4.2GHz Intel(R) Core(TM) i7-7700K CPU and 64 GB of RAM, which we refer to as “desktop” and (ii) a cluster node with 2.5GHz AMD Opteron(TM) Processor 6380 CPU, which we refer to as “HPC cluster” (high performance computing cluster).

We assessed the accuracy of predicted cluster labels from *mbkmeans* and *k*-means using two performance metrics: adjusted Rand index (ARI) [35] and within clusters sum of squares (WCSS). We reasoned that WCSS is a fair performance metric to use as it is the actual objective criterion that is minimized in the *mbkmeans* and *k*-means algorithms and it does not require a “ground truth” label. In contrast, ARI is a metric that assesses the similarity of an estimated cluster label to a known ground truth label.

### Simulated gene expression data

We simulated gene expression data *Y*_*i,g*_ with *i* ∈ (1, …, *N*) cells (or observations) and *g* ∈ (1, … *G*) genes (or features) representing normalized gene expression data in the following way:

i. Assume there are *k* true clusters with probabilities *p*_1_, …, *p*_*k*_. We randomly sample (without replacement) the cluster label for *i*^*th*^ observation *Z*_*i*_ ∈ (1, …, *k*) with probabilities *p*_1_, …, *p*_*k*_.
ii. For the *i*^*th*^ observation with the *k*^*th*^ cluster label, we assume the true biological structure is sampled from a bivariate normal distribution *X*_*ik*_ ∼ *N*_2_(***µ***_*k*_, **Σ**_*k*_) where ***µ***_*k*_ = (*µ*_1*k*_, *µ*_2*k*_), 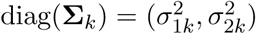 and zero in off diagonals. We combine the *N* observations into a (*N*, 2)-dimensional matrix **X**_(*N*,2)_, which is made up of a mixture of *k* bivariate normal distributions.
iii. Next, we simulate data from a normal distribution **Z**_(*G*,2)_ ∼ *N* (0, 1), to project the *N* observations currently in a 2-dimensional space into a *G*-dimensional space to mimic high-dimensional gene expression data, or **A**_(*N,G*)_ = **XZ**^*T*^.
iv. Finally, we add random noise ***ε***_(*N,G*)_ ∼ *N* (0, 1) to obtain the simulated normalized gene expression data **Y**_(*N,G*)_ = (**Y**_1_, …, **Y**_*N*_), or **Y** = **A** + ***ε***.

### A scRNA-seq experiment with 1.3 million mouse brain cells

A scRNA-seq experiment was performed with the 10X Chromium Genomics platform [25] measuring the gene expression in mouse cells that came from three regions of the brain (cortex, hippocampus, and subventricular zone) and two mouse embryos (E18 C57BL/6 mice). This resulted in a dataset containing *G* = 27,998 genes and *N* = 1,306,127 cells. The data can be downloaded as a sparse HDF5 file directly from the 10X website (https://support.10xgenomics.com/single-cell-gene-expression/datasets/1.3.0/1M_neurons).

However, we used the *TENxBrainData* Bioconductor data package [36], which stores the data as a dense matrix in a HDF5 file. The package returns an object of the `SingleCellExperiment` class [5]. We calculated quality control metrics using the `calculateQCMetrics` function from the *scater* Bioconductor package [37]. We then applied a cell-filter to remove cells with a high proportion of mitochondrial reads (at least 3 median absolute deviations away from the median). We also applied a gene-filter to keep only the genes with at least 1 UMI in at least 1% of the cells. This filtering procedure resulted in a final matrix of 11,720 genes and 1,232,055 cells. Finally, we subsampled cells (random sampling without replacement) and saved each downsampled and pre-processed data as a new HDF5-based `SingleCellExperiment` object. We used all 11,720 genes for the full analysis, while we focused on the 5,000 most variable genes for the subsampling analysis. Since we need fast access to the columns (cells) of the matrix, we saved the data with chunks of dimension (1 × *G*), (i.e., each chunk contained all gene expression measurements for one cell) with default compression level. Unless noted otherwise, we used this chunk geometry to perform all the analyses. In Figure 4, we explore the impact of the HDF5 chunk geometry on the performance of *mbkmeans*.

## Results

One of the main purposes of unsupervised clustering algorithms for the analysis of scRNA-seq data is to empirically define groups of cells with similar expression profiles [5]. We explored the impact of the number of cells and batch sizes in a scRNA-seq dataset on the speed, memory-usage, and accuracy of *mbkmeans* as compared to *k*-means when predicting cluster labels. Specifically, we evaluated the performance of (1) the standard (in-memory) *k*-means algorithm, as implemented in R [8], (2) mini-batch *k*-means applied to in-memory data, (3) and mini-batch *k*-means applied to an on-disk data representation (HDF5), where the last two implementations of mini-batch *k*-means are those available in our *mbkmeans* package. The results reported in the main text are based on a desktop, but results for a HPC cluster are in Supporting Information.

### *mbkmeans* is fast and memory-efficient

Using downsampled datasets ranging from 75,000 to 1,000,000 observations from the 1.3 million mouse brain cells, we found that our on-disk (HDF5) *mbkmeans* uses dramatically less memory than either *k*-means or the in-memory *mbkmeans* for large scRNA-seq datasets (Fig. 1A, Table S1). There is almost no memory increase for datasets with larger sample sizes using our on-disk implementation of *mbkmeans*, and we can cluster 1 million cells with only 1.55GB of RAM, as compared to 39.4 GB for the in-memory version. We were not able to compare to standard *k*-means at large sample sizes due to lack of sufficient memory; however, when using 300,000 cells, *k*-means used 52 GB as opposed to 0.98GB with the on-disk *mbkmeans*. In addition, the in-memory *mbkmeans* uses far less memory than *k*-means, requiring only 11.95GB for 300,000 cells.

**Figure 1:**
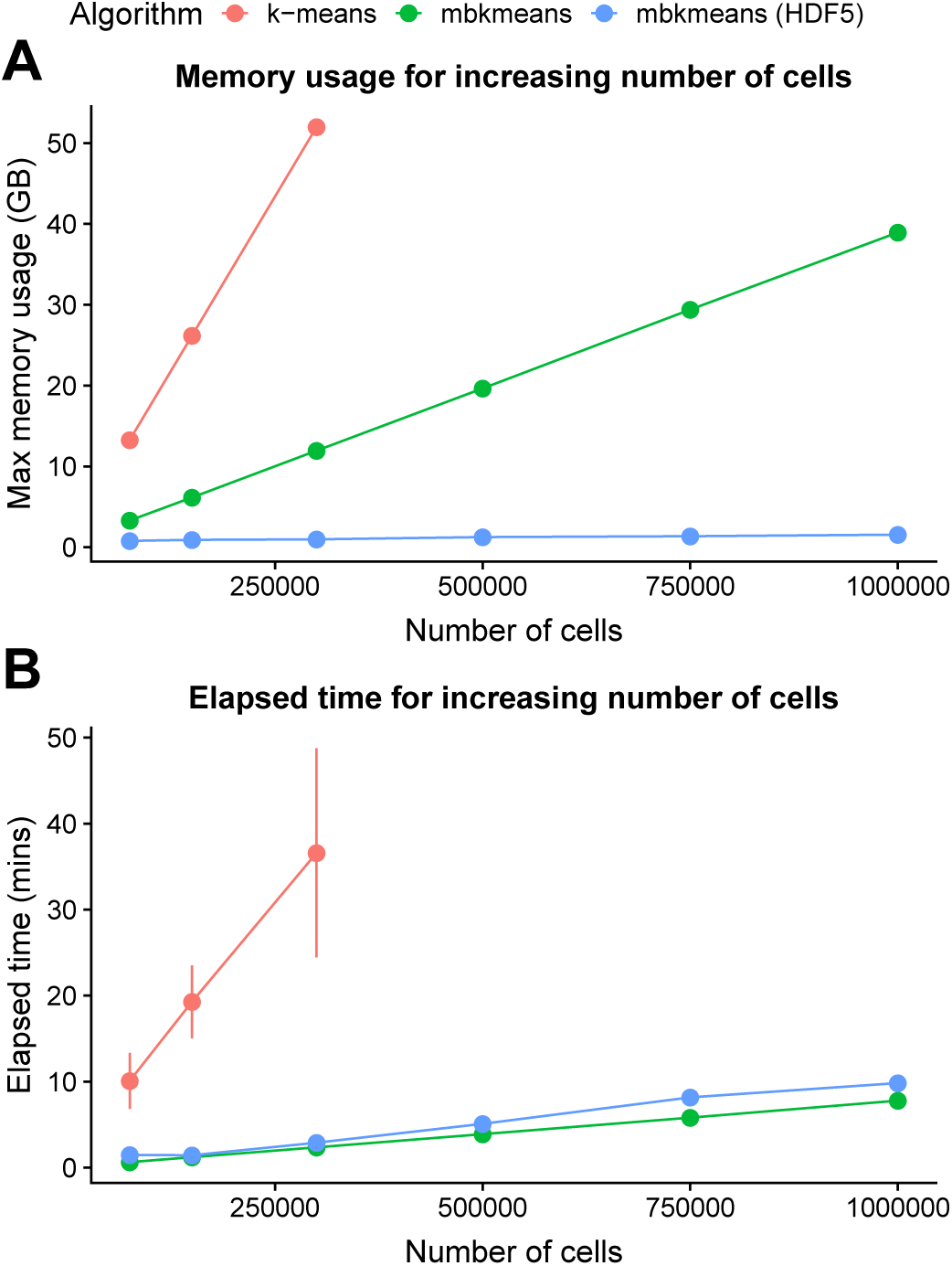
*mbkmeans* uses less memory and is faster than *k*-means. Performance evaluation (y-axis) of **(A)** maximum memory (RAM) used (GB) and **(B)** elapsed time (minutes) (repeated 10 times) for increasing sizes of datasets (x-axis) with *N* = 75,000, 150,000, 300,000, 500,000, 750,000, and 1,000,000 observations and *G* = 5,000 genes, using a desktop. Results for *mbkmeans* are in blue (in-memory) and green (on-disk); *k*-means is in red. We used *k* = 15 for both algorithms and used a batch size of *b* = 500 observations for *mbkmeans*.

Furthermore, we found both of our *mbkmeans* implementations are significantly faster compared to *k* - means (Fig. 1B, Table S1). Specifically, we can cluster 1 million cells in 9.8 and 7.8 minutes (mean values across 10 runs) for in-memory and on-disk implementations of mini-batch *k*-means, respectively, compared to 36.6 minutes using in-memory *k*-means for 300,000 cells (*k*-means fails to complete with larger datasets).

While the above results were using a desktop computing system, the results from the HPC cluster were nearly identical in memory-usage, though all of the algorithms take slightly longer (Fig. S1, Fig. S2).

### *mbkmeans* is accurate

Using simulated gene expression data, we explored the effect of increasing sizes of datasets, as well as batch sizes, on the performance of the algorithms. We found that using datasets with a batch size *b* = 500 observations or larger led to no loss in accuracy with respect to *k*-means, based on ARI (Fig. 2A, Table S2) and WCSS (Fig. 2B, Table S2). In addition, we considered a wider range of sizes of datasets and found consistent ARI and WCSS results using both a desktop (Fig. S3, Fig. S4) and HPC Cluster (Fig. S5, Fig. S6).

**Figure 2:**
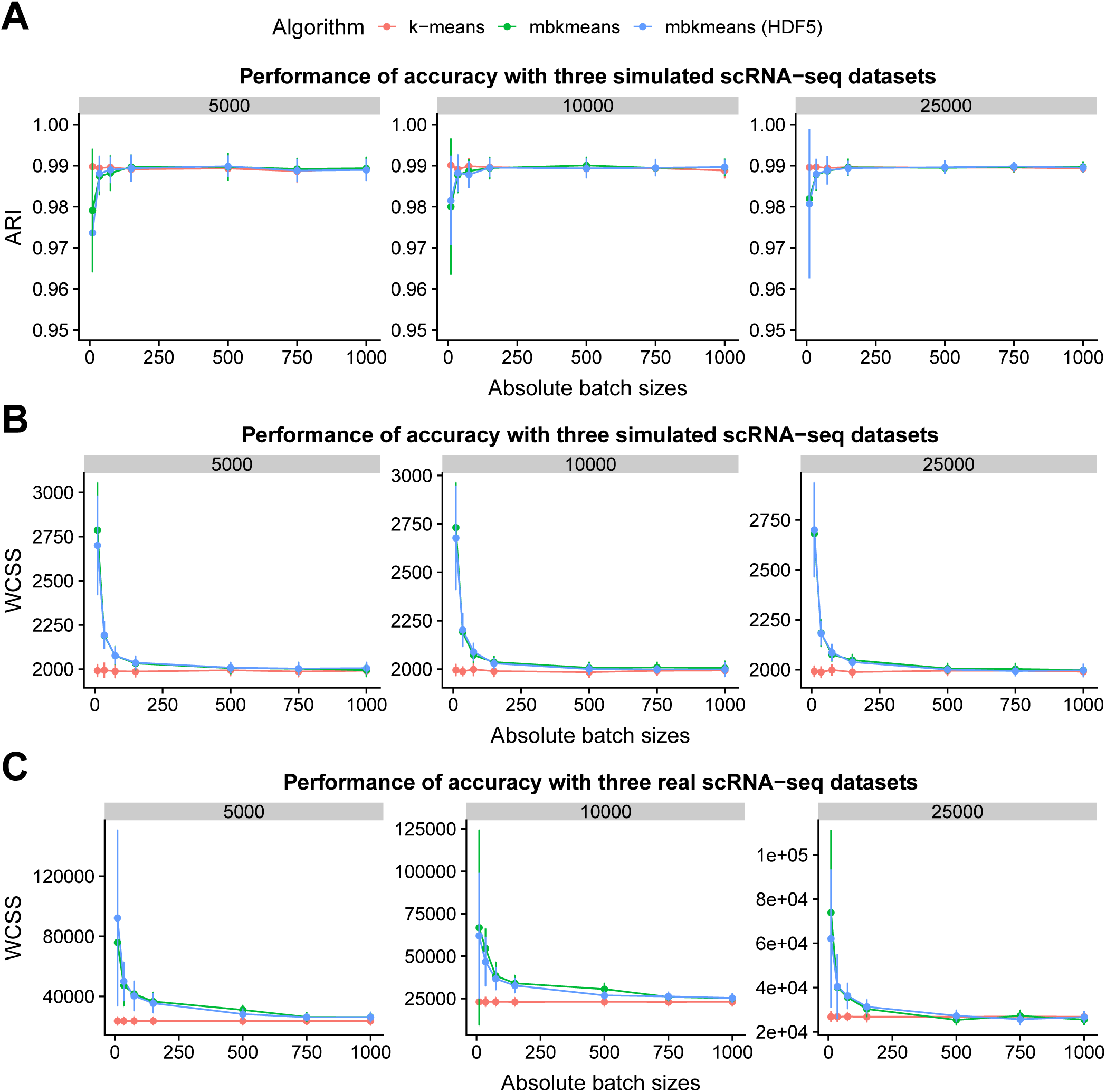
The accuracy of *mbkmeans* depends on batch size. Performance evaluation (y-axis) with **(A)** adjusted Rand index (ARI) and **(B)** within clusters sum of squares (WCSS) for increasing batch sizes ranging from 75 to 1000 cells (x-axis) using simulated gene expression data (*G* = 1000) with a fixed *k* = 3 true centroids with three sizes of datasets (*N* = 5000, 10000, 25000). **(C)** WCSS (y-axis) for increasing batch sizes (x-axis) using real scRNA-seq gene expression data from 10X Genomics and *k* = 15 for both algorithms. ARI and WCSS is reported as an average across 50 runs.

Next, we assessed the WCSS using the 1.3 million mouse brain cells by downsampling to similar dataset sizes (*N* = 5,000, 10,000, and 25,000). We found similar results to simulated data (Fig. 2C, Table S2) with no substantial difference in the minimum observed WCSS between the *k*-means and *mbkmeans* algorithms (using *k* = 15) if using a batch size of *b* = 500 observations or more. These results demonstrate that, as long as the batch size of the *mbkmeans* is not unreasonably small (at least 500 − 1, 000 cells in a batch), the *mbkmeans* algorithm is as accurate as the standard *k*-means algorithm.

Finally, because one of the user-defined parameters in the *k*-means and mini-batch *k*-means algorithms is the number of centroids (*k*), we investigated the impact of *k* on both memory-usage and accuracy. We found that the maximum memory (GB) used was not affected by *k* either in the in-memory or on-disk implementations for small (*N* = 25,000) or large (*N* =1,000,000) datasets in both a desktop or HPC cluster (Fig. S7). However, using simulated scRNA-seq data we found that accuracy – using ARI (Fig. S8) or using WCSS (Fig. S9, Fig. S9) – varied as a function of *k* with the highest accuracy resulting when *k* = 15 (the true simulated centroids), as expected.

### Impact of *mbkmeans* batch size on speed and memory-usage

The choice of batch size (*b*) in *mbkmeans* can in principle have an impact on how long it takes the algorithm to run and how much memory is used. We used the downsampled 1.3 million mouse brain cells dataset with *N* = 1,000,000 observations and considered the effect of increasing batch sizes for both the in-memory and on-disk implementations of *mbkmeans*. We found that there was no impact on maximum memory usage (Fig. 3A, Table S3) nor speed (Fig. 3B, Table S3) for batch sizes up to *b* = 10,000. This is perhaps unsurprising, since computations in memory on datasets of size 10,000 are routine and unlikely to have noticeable differences in time and memory. For larger batch sizes, we started to see noticeable increases in demand for memory and larger time to run *mbkmeans*. The memory-usage results were consistent across a desktop and HPC cluster (Fig. S10), but took longer to complete on a HPC cluster (Fig. S11).

**Figure 3:**
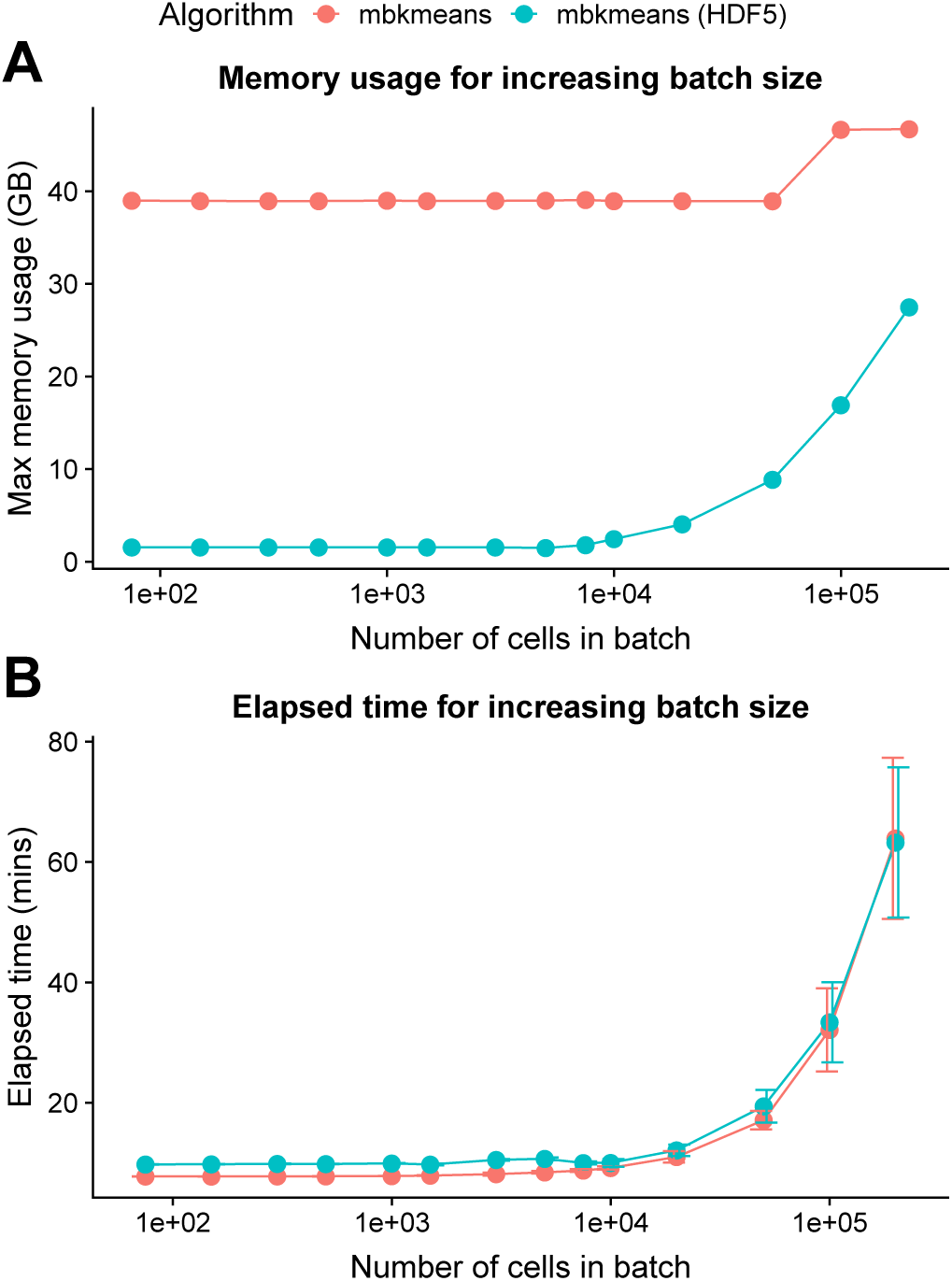
The speed and memory-usage of *mbkmeans* depends on batch size. Performance evaluation (y-axis) of **(A)** maximum memory (RAM) used (GB) and **(B)** elapsed time (minutes) for increasing batch sizes (x-axis) with *b* = 75, 150, 300, 500, 1,000, 1,500, 3,000, 5,000, 7,500, 10,000, 20,000, 50,000, 100,000, and 200,000 with a dataset of size *N* = 1,000,000 observations using a desktop. Results for *mbkmeans* in-memory are in red and and on-disk in blue. We used *k* = 15 for the number of centroids.

Combined with the results of accuracy, these results demonstrated that the algorithm is not sensitive to the batch size parameter, with values in the range of *b* = 500 to 10,000 giving comparable results with respect to both accuracy and computational performance.

### Impact of HDF5 file format geometry can improve performance

How the data are stored (or ‘chunked’) and accessed in the HDF5 format (or different file geometries) can affect the speed of accessing the data. For example, the indexing can be based on columns or rows of the data (vertical or horizontal slices), or other sub-matrices (rectangles). In the case of a matrix (two-dimensional array), the default chunk size in the *HDF5Array* package selects a rectangle that satisfies the following constraints: (i) its area is less than 1,000,000; (ii) it fits in the original matrix; (iii) its shape is as close as possible as the shape of the original matrix (the two dimensions are in the same proportion).

We investigated the choice of different file chunking geometries on the performance of the *mbkmeans* algorithm, and we found that the best choice for minimizing the maximum RAM used is to index a HDF5 file by the cells (or observations) of the matrix (Fig 4A, Table S4). In the application of scRNA-seq with a `SingleCellExperiment` object, this corresponds to indexing by columns. In contrast, if a file is indexed by genes (or rows in a `SingleCellExperiment` object), we found the on-disk *mbkmeans* implementation required twice as much RAM – though even then it is still a relatively small memory footprint compared to an in-memory version. Similarly, we found that indexing by cells also improves the speed of the algorithm (Fig 4B, Table S4). We also considered indexing a HDF5 file by the entire matrix in one chunk (“single chunk” in Fig 4), which is the default in the HDF5 library format [24]. In this case, we found that even for a small number of cells, there is an substantial memory cost for this naive geometry. In addition, in all of the above scenarios, we found similar results using a HPC cluster for both memory-usage (Fig. S12) and speed (Fig. S13).

**Figure 4:**
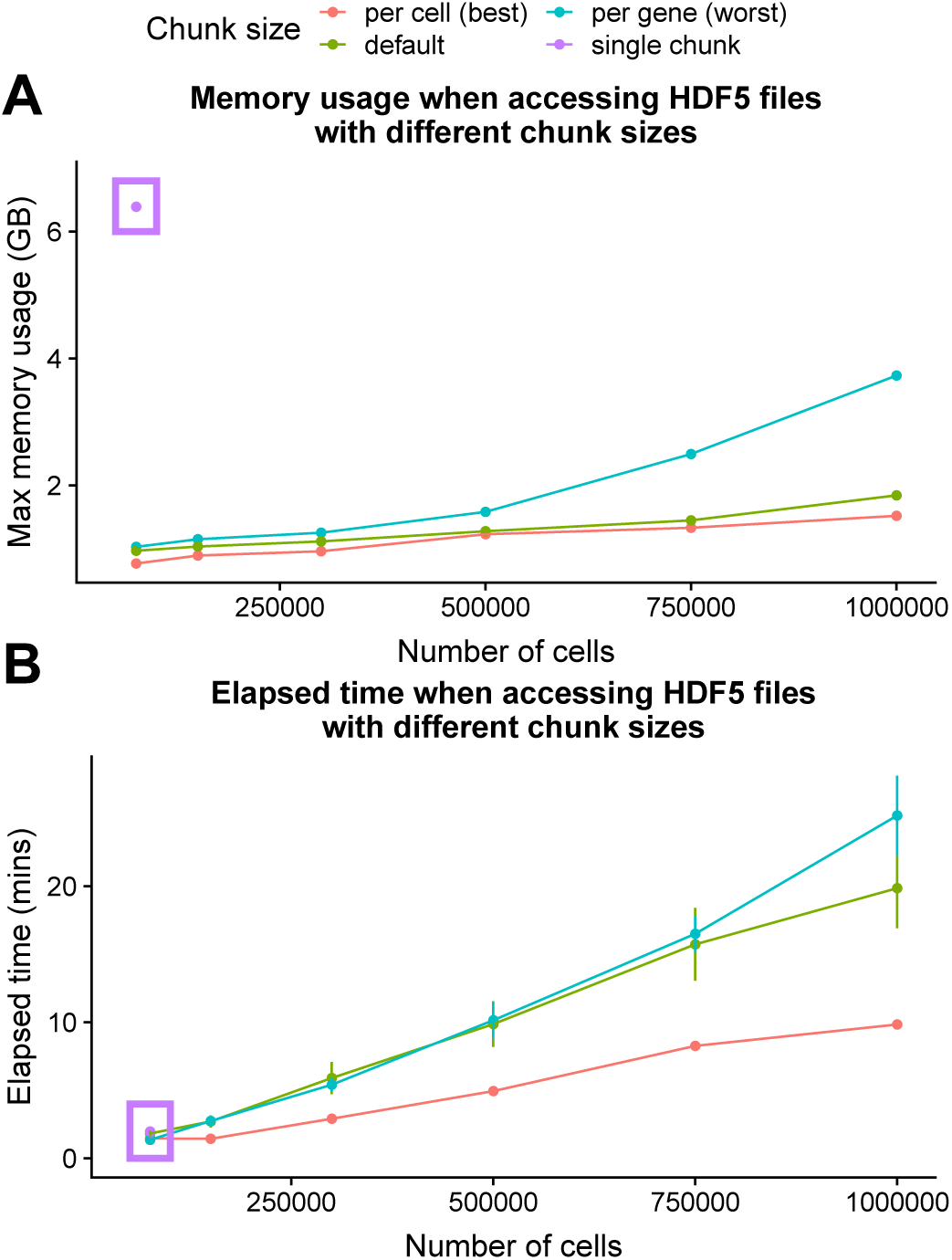
The speed and memory-usage of the on-disk *mbkmeans* implementation depends on the structure of the on-disk file. Performance evaluation (y-axis) of **(A)** maximum memory (RAM) used (GB) and **(B)** elapsed time (minutes) (repeated 10 times) for increasing sizes of datasets (x-axis) with *N* = 75,000, 150,000, 300,000, 500,000, 750,000, and 1,000,000 observations using a desktop. Results for indexing a HDF5 file by gene is blue, by cell is red, as a single chunk is purple and the default indexing is green. The single chunk was only able to run for the smallest dataset size (*N* = 75,000). We used *k* = 15 and used a batch size of *b* = 500 observations for *mbkmeans*.

Finally, we note that while the default geometry implemented in *HDF5Array* is not optimal for clustering, there are many different components in a standard scRNA-seq pipeline, with clustering typically not being the slowest step (see ‘A complete analysis of a large single-cell dataset’ Section below), and thus the choice of geometry may ultimately be better determined by the performance of these other steps.

### A complete analysis of a large single-cell dataset

We analyzed the full 1.3 million mouse brain cells with all the steps in a standard scRNA-seq analysis [5]: quality control and filtering, normalization, dimensionality reduction using Principal Components Analysis (PCA), and finally clustering for detection of subtypes. For all of the non-clustering steps, we used recent packages in Bioconductor that operate directly on HDF5 files. Specifically, we used *scater* [37] *for quality control and dimensionality reduction, via the BiocSingular* package, which implements the implicitly restarted Lanczos bidiagonalization algorithm (IRLBA) [38], and *scran* [39] for normalization. We note that we used *mbkmeans* as a preliminary step to create homogeneous cell groups for *scran* normalization (see [39] for details). Our *mbkmeans* package complements these existing packages, allowing us to demonstrate a complete HDF5-ready pipeline for scRNA-seq analysis in R/Bioconductor.

We note that because clustering is generally performed after a dimensionality reduction (e.g. reducing to the top 50 principal components or PCs), the size of the data matrix to be clustered is significantly reduced in complexity and can be potentially performed in-memory, even with 1.3 million cells. However, some normalization methods, such as *scran*, make use of an initial clustering to improve the accuracy of the normalization, which is before the dimensionality reduction step; in this case the dimension of the clustering problem is over all cells and all expressed genes and benefits greatly from using on-disk clustering.

In Table S5, we show the compute time for each step of the pipeline. The clustering of the data with *mbkmeans*, even with all genes, was a very quick part of the pipeline, taking roughly 9 minutes (with *k* = 15 and batch size of 500); in contrast, normalization took 5 hours, and IRLBA PCA took 96 hours. On the computationally simpler problem of clustering on only the top 50 PC dimensions, which we performed via the in-memory *mbkmeans*, the clustering took only about 30 seconds – quickly enough to rerun the clustering algorithm with different number of clusters or on random subsamples of the data for stability analysis.

In Fig. 5A we show the UMAP of the full dataset, with cells color-coded by their assignment into the 15 final clusters found by applying *mbkmeans* on the first 50 PCs. *mbkmeans* clearly identified distinct outlying groups of cells, such as clusters 2, 5, 12, 14, and 15 (Table S6). The remaining clusters are not easily separated by eye in two dimensions, making it difficult to discern whether *mbkmeans* is identifying meaningful clusters. To further evaluate the clusters, we accumulated a list of genes previously shown to discriminate known subtypes of cells in developing and adult mouse brains [40, 41, 42, 43, 44, 45, 46]. We show the average expression of each cluster for these marker genes in Fig 5B. Many of the clusters identified by *mbkmeans* have unique expression of these marker genes, indicating that *mbkmeans* is finding meaningful biological clusters. For example, clusters 1, 7, and 10 – the boundaries of which are not obviously distinguished by eye on the UMAP representation – all correspond to Radial Precursors [44], but they each have clear markers that distinguish them, suggesting that they correspond to different developmental stages (see Table S6). Similarly, cluster 8 (expressing the L2/3 marker *Crym* [41]) and 11 (expressing the L5/6 marker *Ntf3* [40]) represent two distinct pyramidal excitatory neuron populations, possibly residing in different cortical layers.

**Figure 5:**
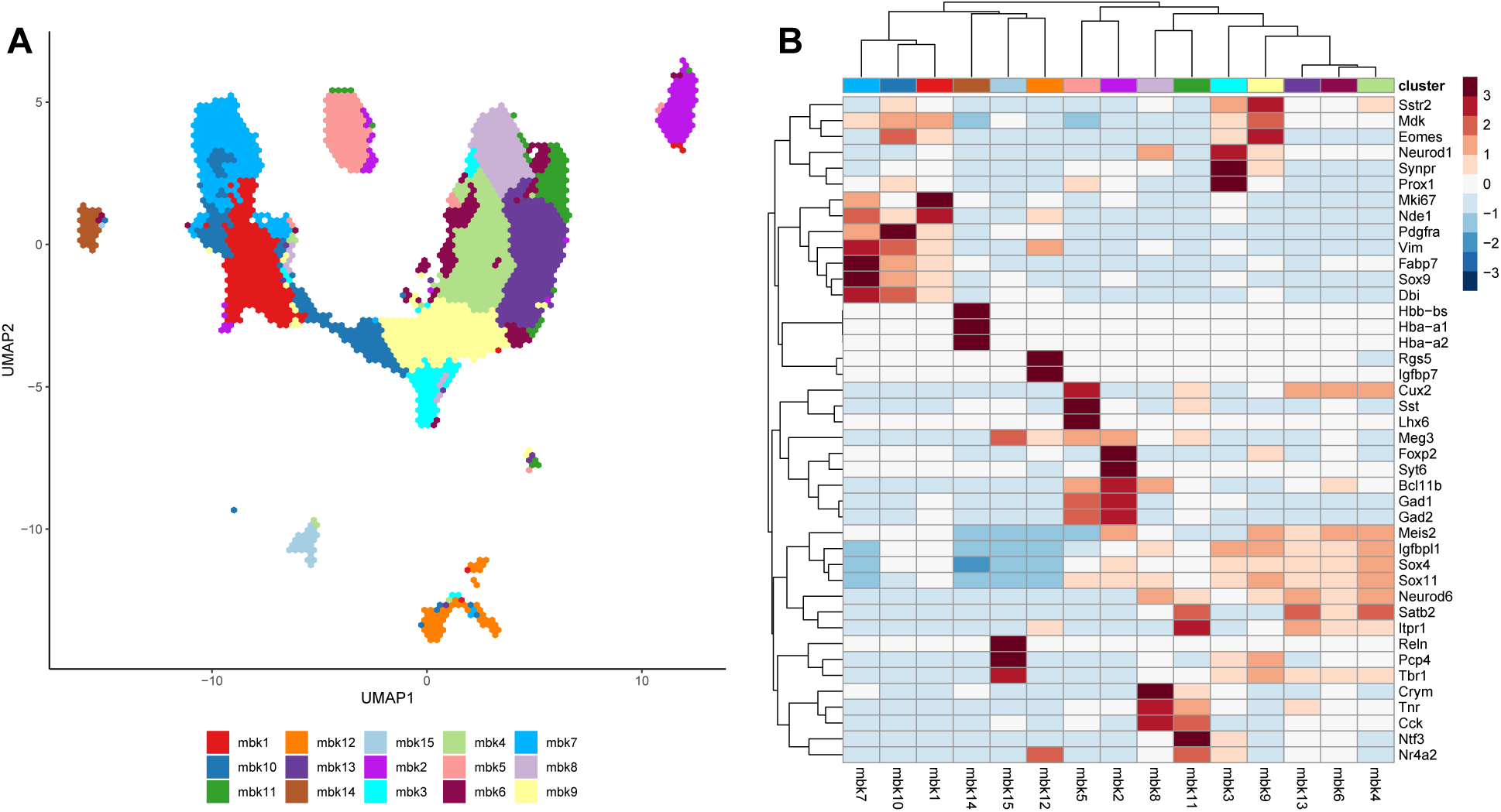
Results of full analysis on 1.3 million mouse brain cells. **(A)** Hexbin plot [47] of the UMAP representation of the 1.3 million cells, color coded by the clusters found via *mbkmeans*. **(B)** Heatmap of the average gene expression of each of the 15 clusters found by *mbkmeans* for 42 marker genes.

## Availability and future directions

A major challenge in the analysis of scRNA-seq data is the scalability of analysis methods as datasets increase in size over time. This is particularly problematic as experiments now frequently produce millions of cells [48, 49, 50, 51], making it challenging to even load the data into memory. Providing analysis methods, such as unsupervised clustering, that do not require data to be loaded into memory is an imperative step for scalable analyses. While large-scale scRNA-seq data are now routinely stored in on-disk data formats (e.g. HDF5 files), the methods to process and analyze these data are lagging.

To address this, we have developed an open-source implementation of the mini-batch *k*-means algorithm to provide an unsupervised clustering algorithm scalable to millions of observations. Unlike other existing implementations of mini-batch *k*-means, our algorithm harnesses the structure of the mini-batch *k*-means algorithm to only read in the data needed for each batch, controlling memory usage for large datasets. This makes our implementation truly scalable and applicable to both standard in-memory matrix objects, including sparse matrix representations, and on-disk data representations that do not require all the data to be loaded into memory at any one time, such as HDF5 matrices. We have demonstrated the performance improvement of the *mbkmeans* package across a range of different sized datasets, both with simulated and real single-cell datasets. We have also benchmarked an end-to-end Bioconductor pipeline, which includes using *mbkmeans* for subtype discovery, on a 1.3 million scRNA-seq dataset.

Our implementation of mini-batch *k*-means is available as the open-source *mbkmeans* package in Bioconductor (https://bioconductor.org/packages/mbkmeans). The analyses in this manuscript were performed using *mbkmeans* version 1.4.0, and all code to replicate the analyses is available at: https://github.com/stephaniehicks/benchmark-hdf5-clustering.

## Acknowledgments

The authors would like to thank Hervé Pagès, Mike Smith, Kasper Hansen, and Pete Hickey for helpful discussions on using HDF5 files in the Bioconductor framework and John Ngai for his help with the cell type assignments.

## Contributions

SCH, EP, DR developed the idea of the project and package. YN and DR created the open-source R/Bioconductor software package. RL, SCH and DR performed the benchmarking data analyses. SCH, EP, and DR wrote the manuscript with contributions and input from all authors. All authors read and approved the final manuscript.

## Supporting information

**Fig. S1.**
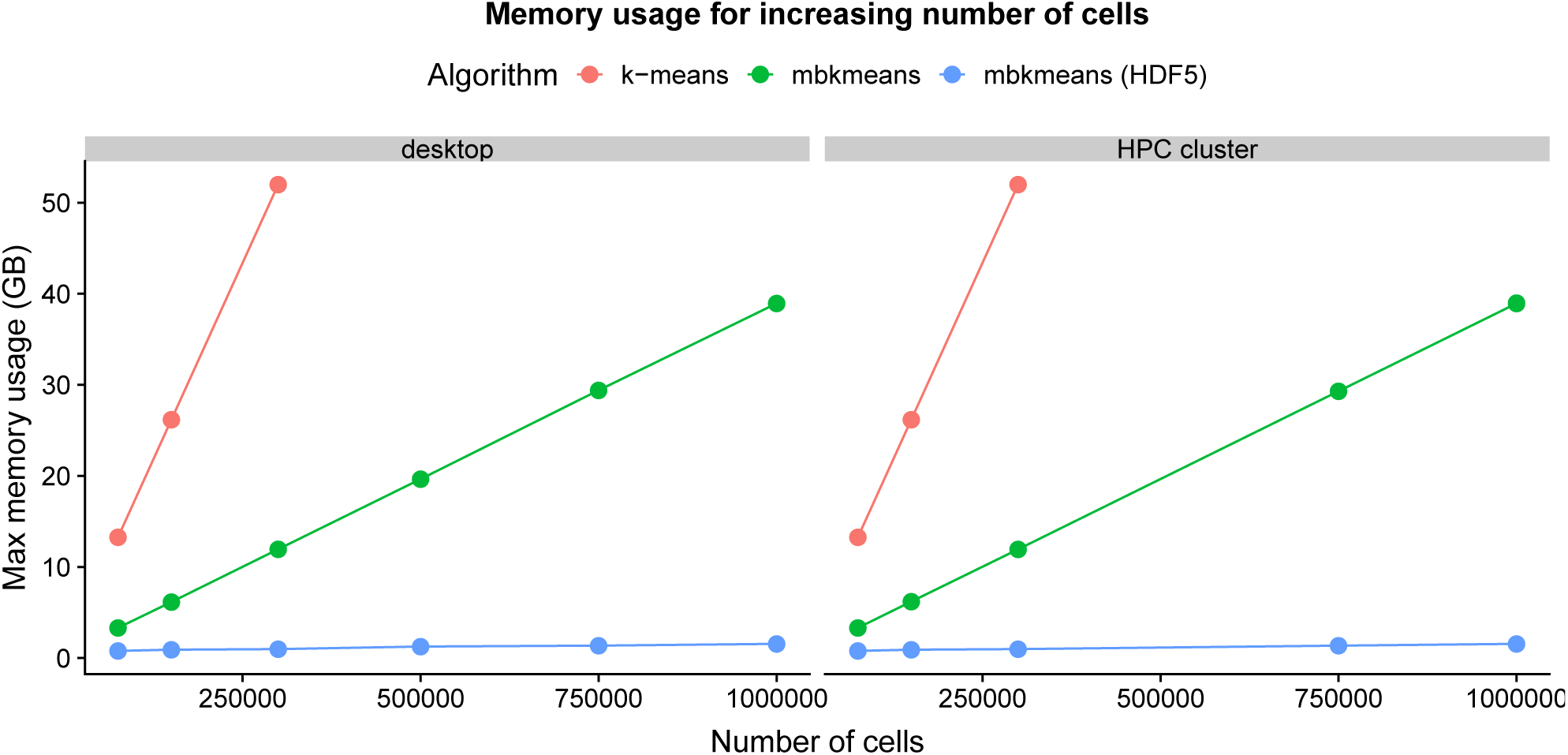
Memory-usage reported for both desktop and HPC cluster corresponding to Fig. 1.

**Fig. S2.**
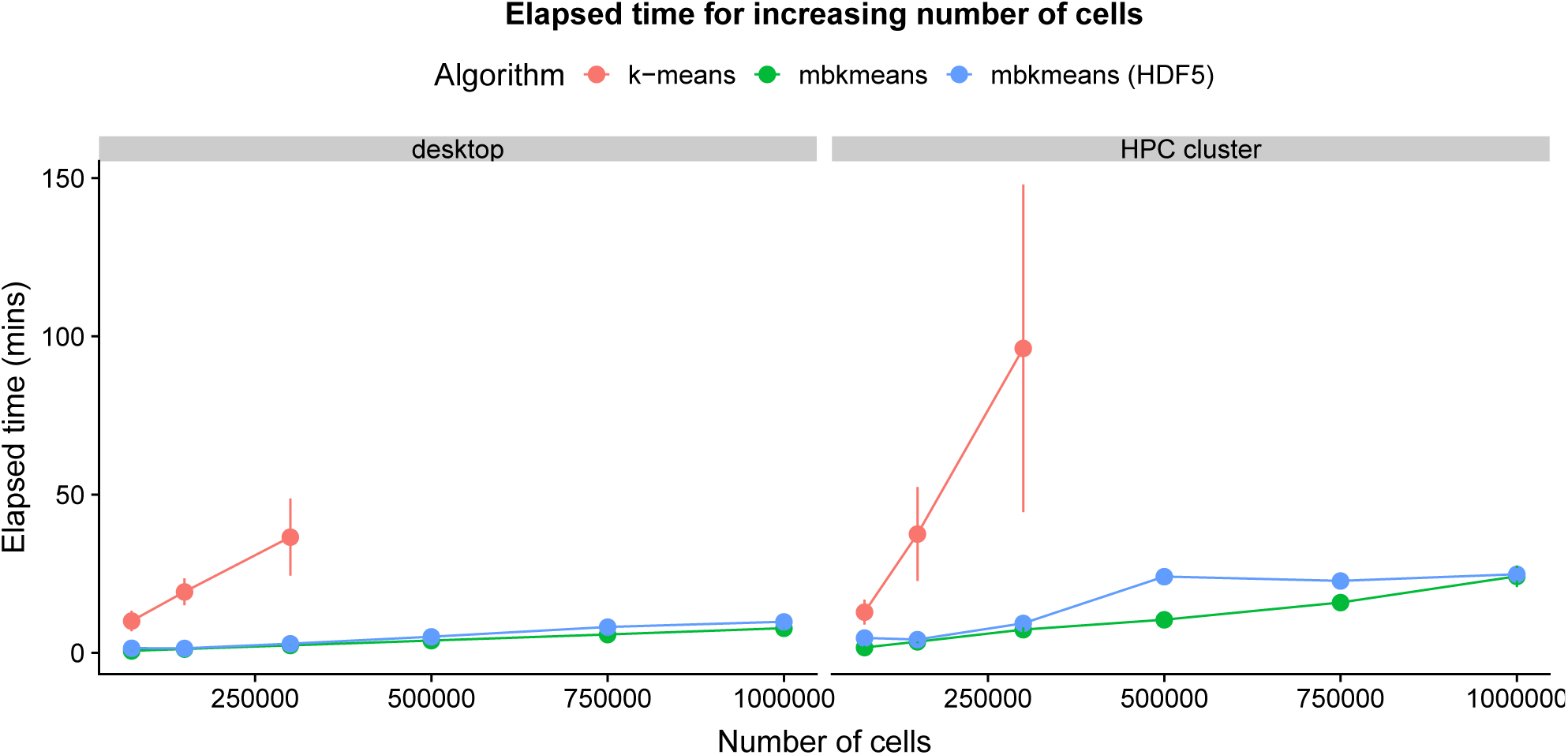
Elapsed time (minutes) reported for both desktop and HPC cluster corresponding to Fig. 1.

**Fig. S3.**
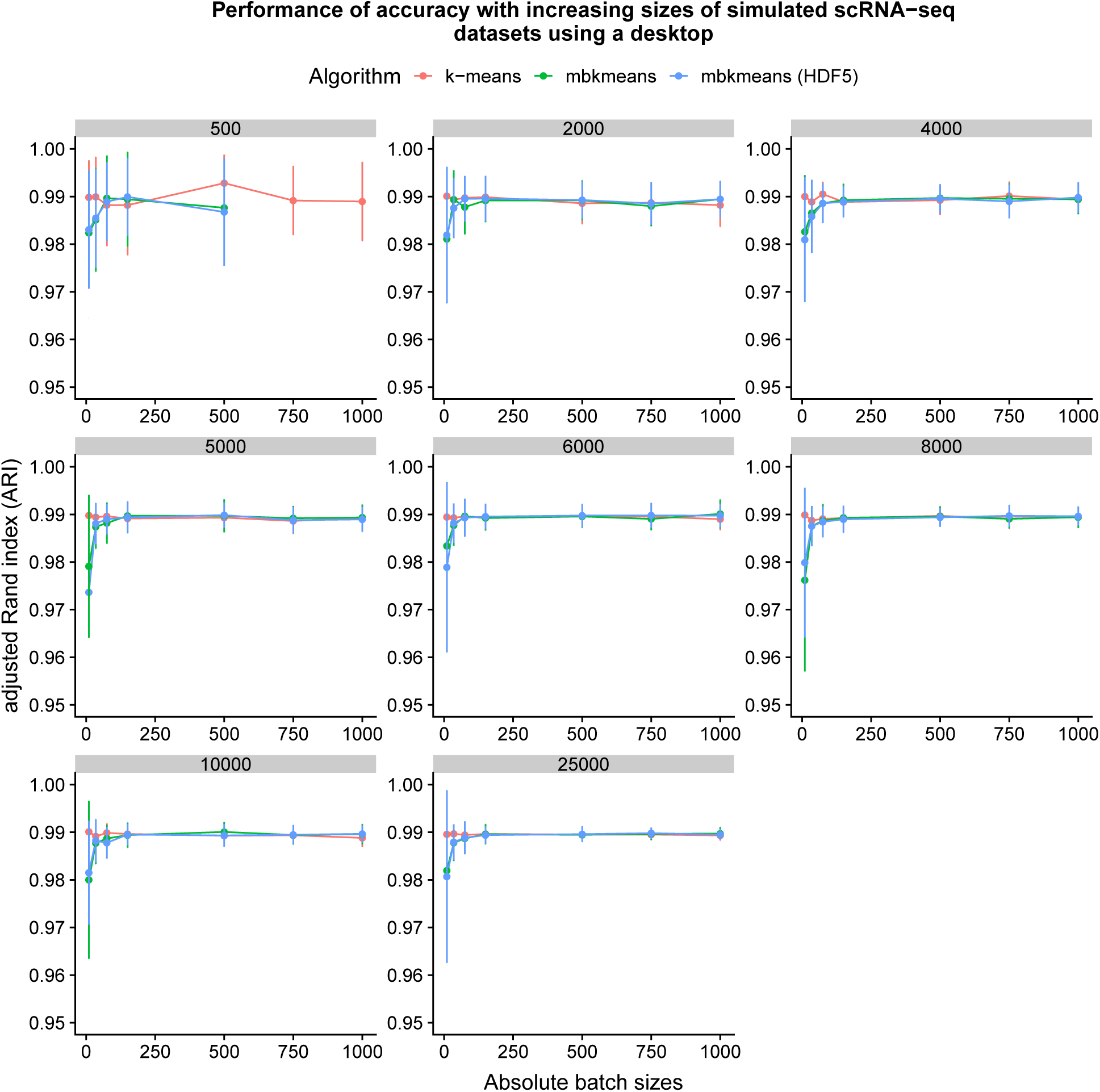
Accuracy (ARI) corresponding to Fig. 2A using simulated data (*N* = 500, 2,000, 4,000, 5,000, 6,000, 8,000, 10,000, 25,000 observations) reported using a desktop.

**Fig. S4.**
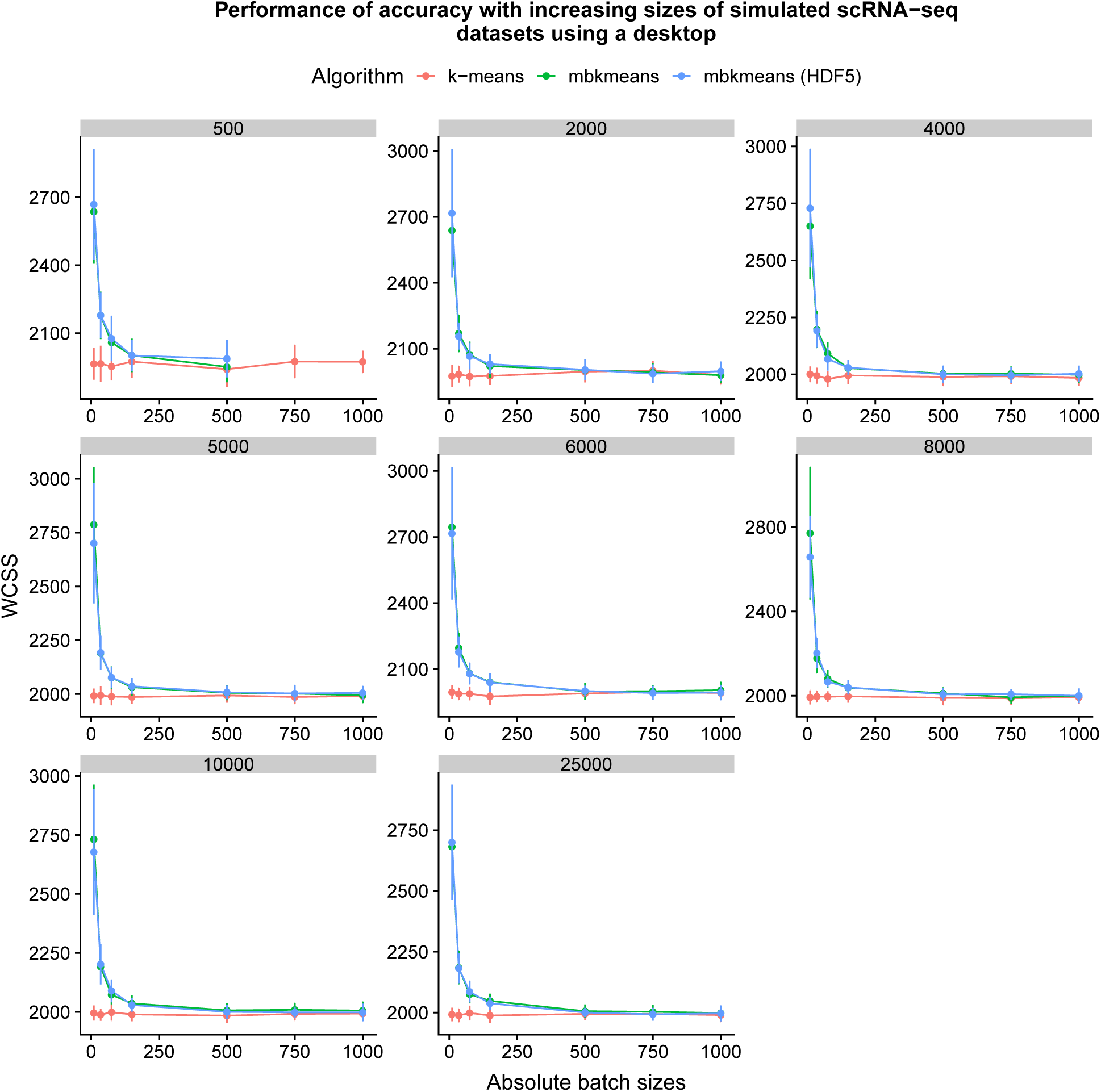
Accuracy (WCSS) corresponding to Fig. 2B using simulated data (*N* = 500, 2,000, 4,000, 5,000, 6,000, 8,000, 10,000, 25,000 observations) reported using a desktop.

**Fig. S5.**
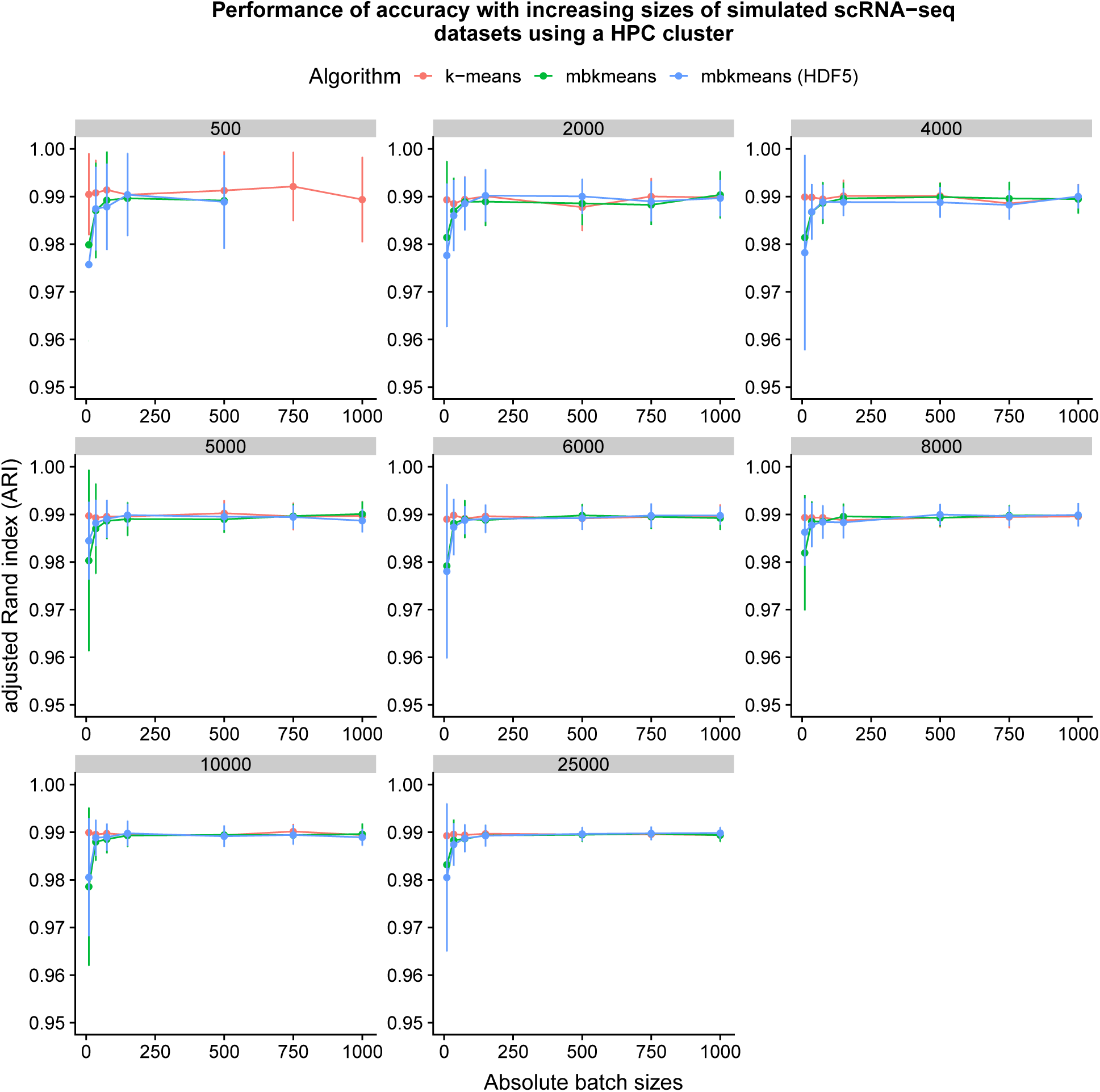
Accuracy (ARI) corresponding to Fig. 2A using simulated data (*N* = 500, 2,000, 4,000, 5,000, 6,000, 8,000, 10,000, 25,000 observations) reported using a HPC cluster.

**Fig. S6.**
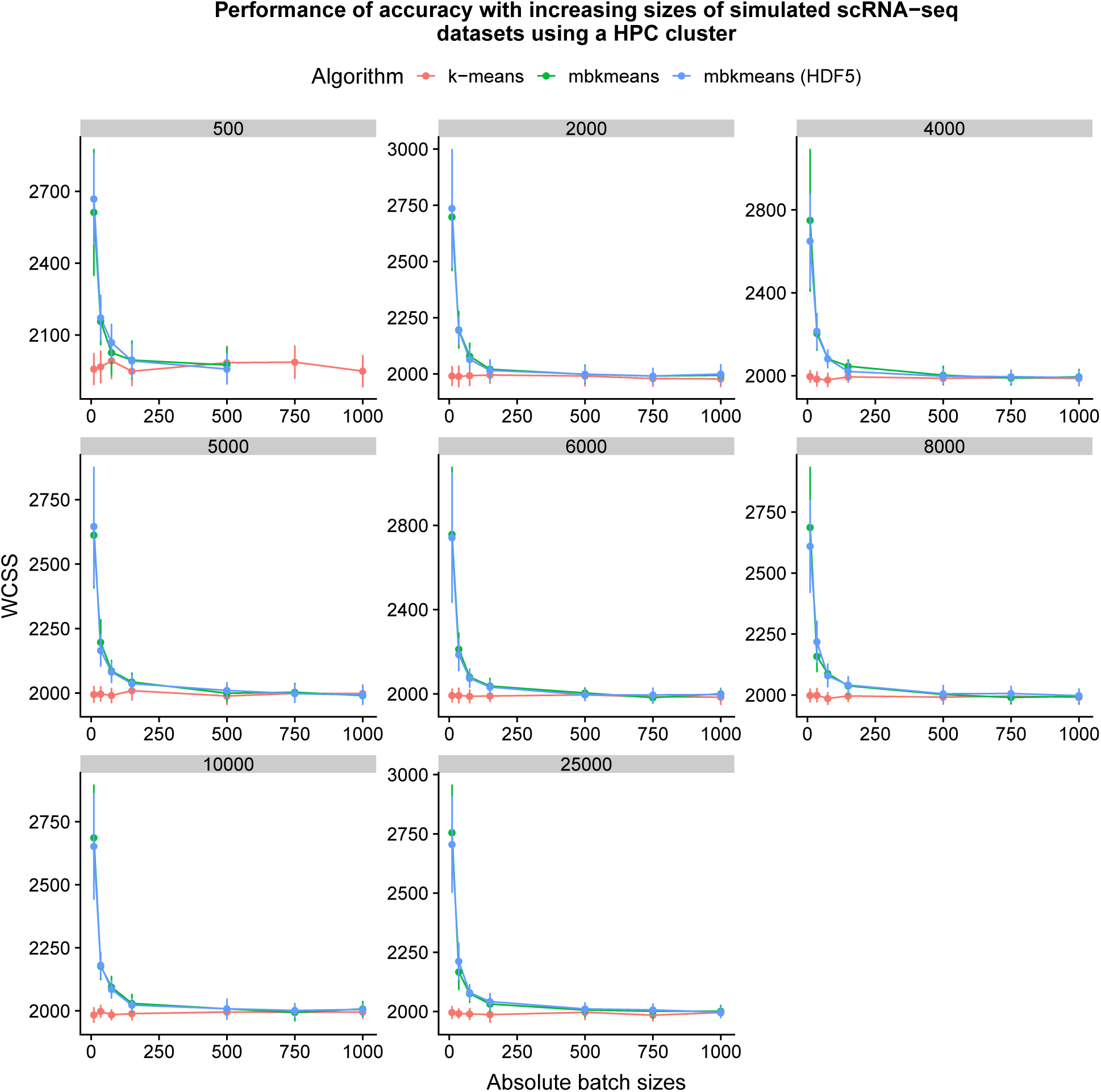
Accuracy (WCSS) corresponding to Fig. 2B using simulated data (*N* = 500, 2,000, 4,000, 5,000, 6,000, 8,000, 10,000, 25,000 observations) reported using a HPC cluster.

**Fig. S7.**
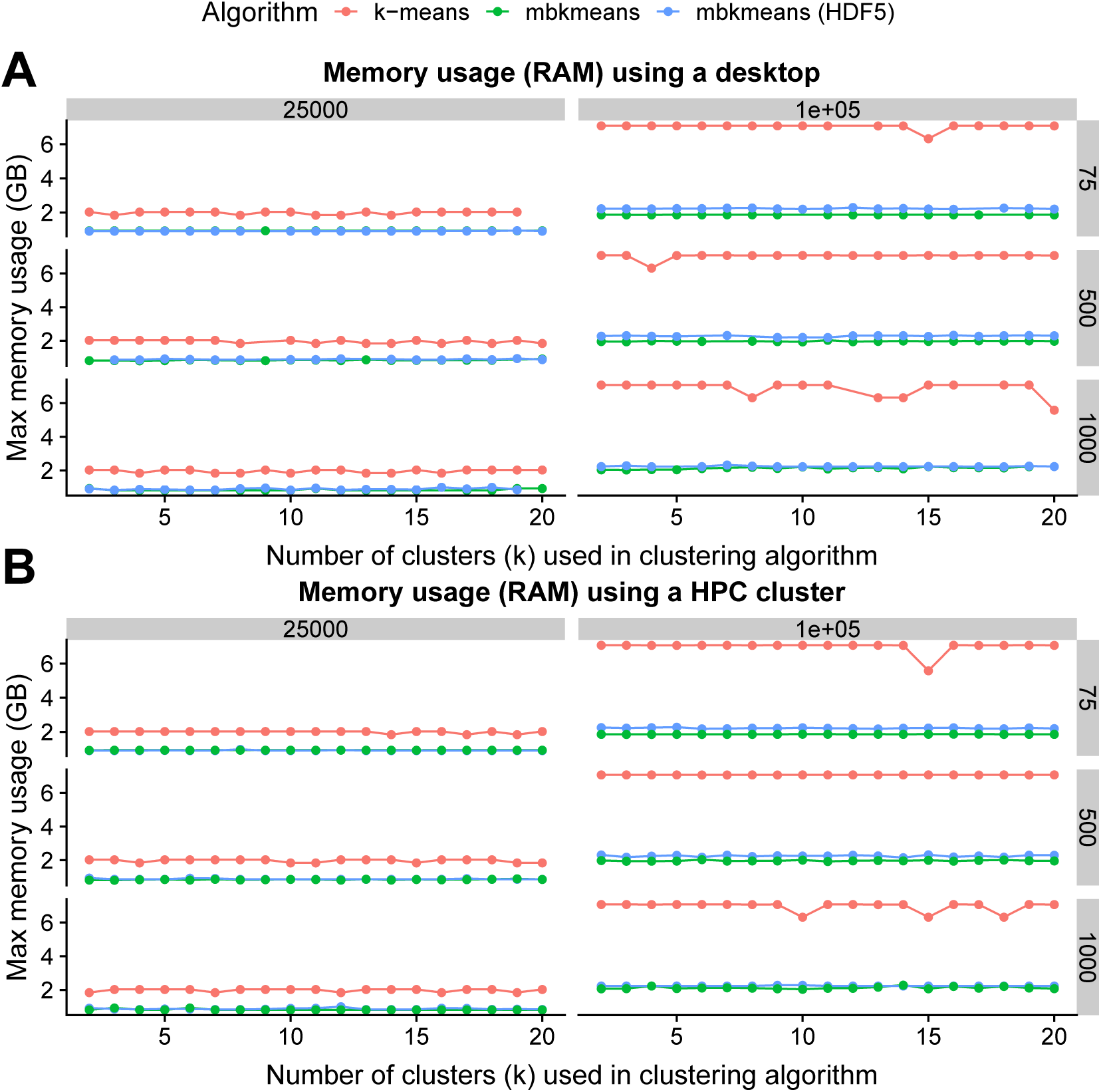
Memory usage with two sizes of simulated scRNA-seq datasets and three absolute batch sizes (75, 500, 1000) with the true number of clusters as k = 15.

**Fig. S8.**
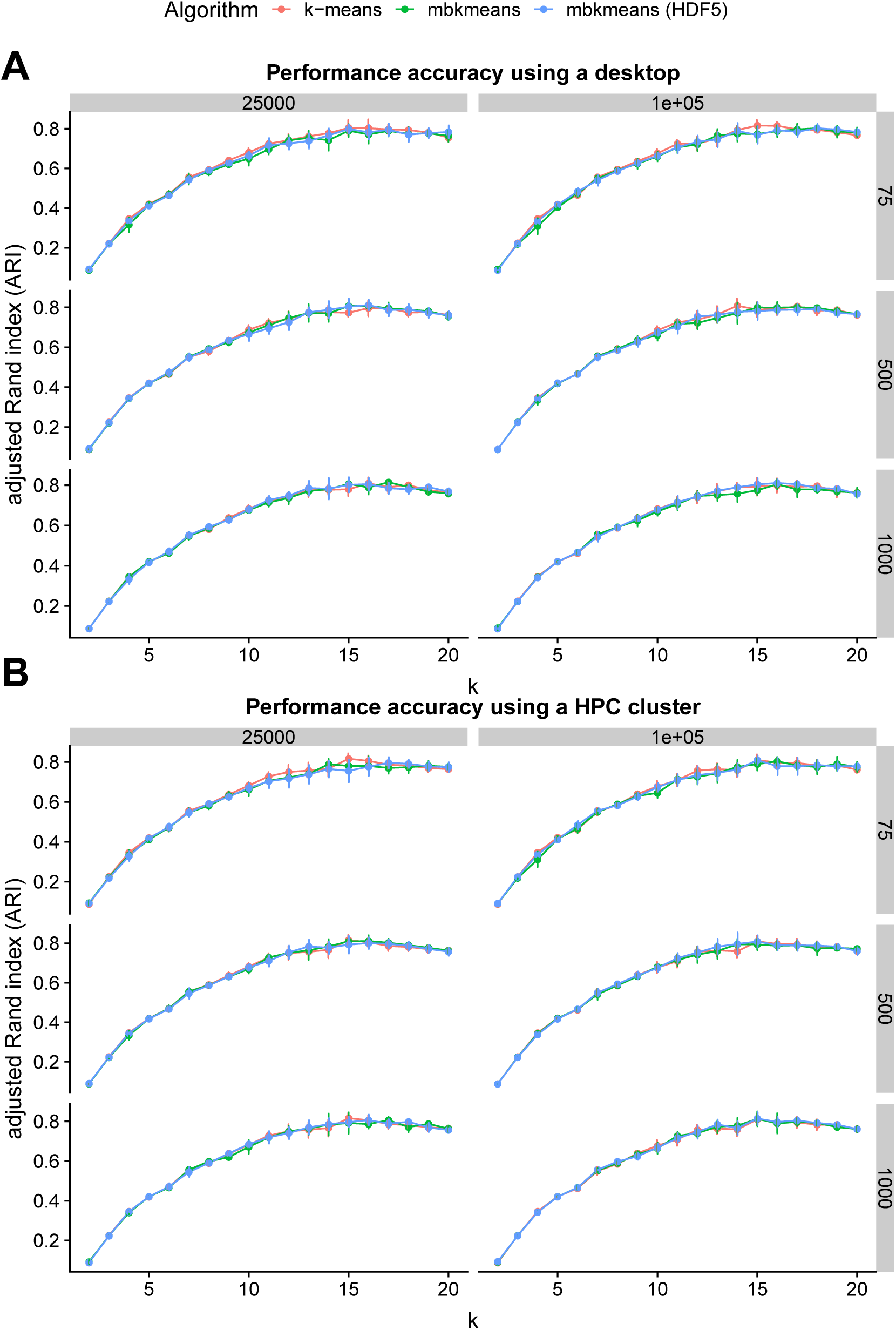
Performance evaluation of ARI with increasing estimated cluster centroids *k* using *mbkmeans* and *k*-means using a desktop and HPC cluster. We simulated gene expression data with 15 true centroids for two sizes of datasets (*N* = 25000, 100000, both using *G*=1000 genes) considered three absolute batch sizes of cells (*b*= 75, 500, 1000) for *mbkmeans* (both in memory and on-disk using HDF5 files using a desktop). We show the impact of increasing the number of estimated cluster centroids *k* used in the clustering algorithm (x-axis) on the adjusted Rand index (ARI) performance metric (y-axis).

**Fig. S9.**
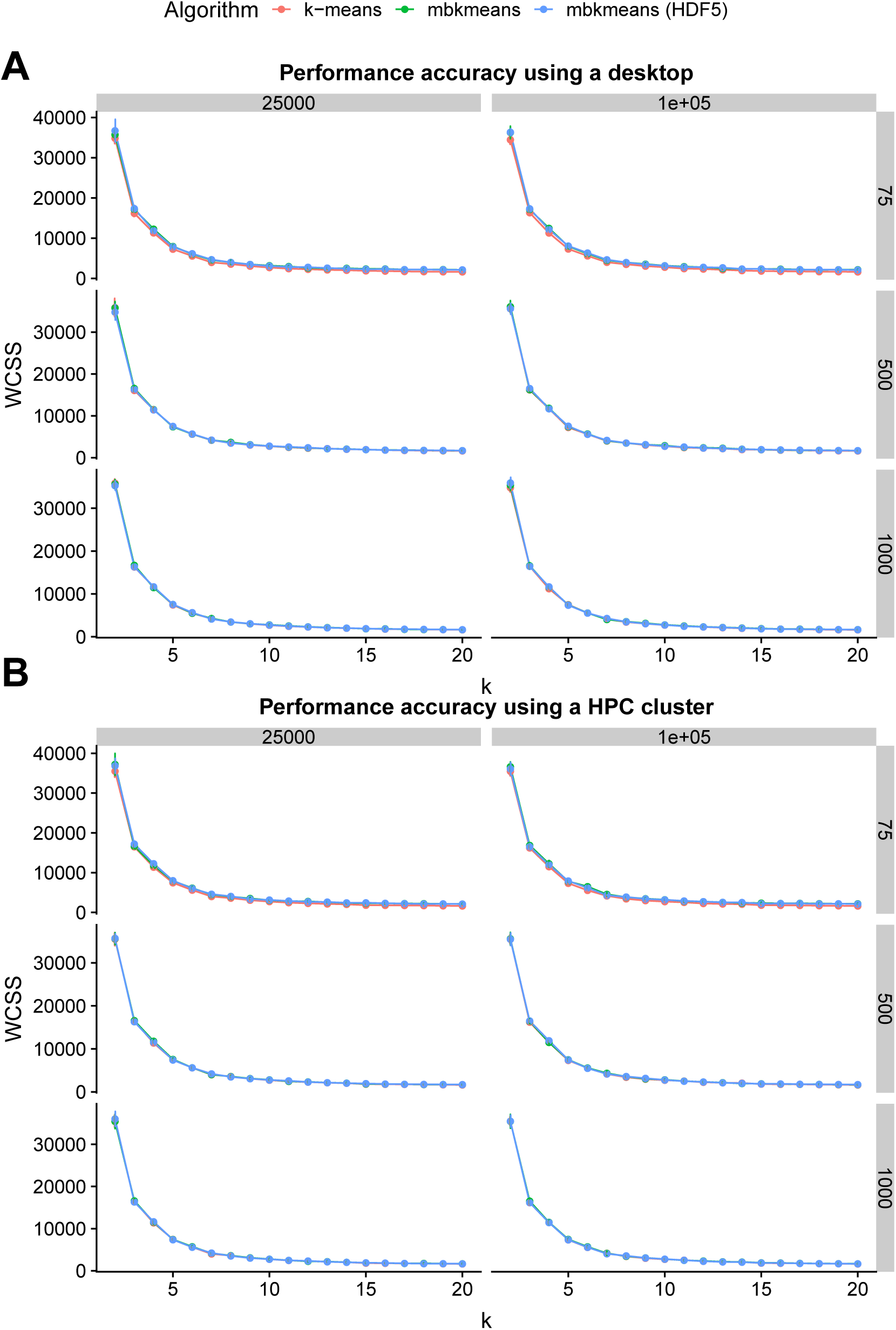
Performance evaluation of WCSS with increasing estimated cluster centroids *k* using *mbkmeans* and *k*-means using a desktop and HPC cluster. We simulated gene expression data with 15 true centroids for two sizes of datasets (*N* = 25000, 100000, both using *G*=1000 genes) considered three absolute batch sizes of cells (*b*= 75, 500, 1000) for *mbkmeans* (both in memory and on-disk using HDF5 files using a desktop). We show the impact of increasing the number of estimated cluster centroids *k* used in the clustering algorithm (x-axis) on the within clusters sum of squares (WCSS) performance metric (y-axis).

**Fig. S10.**
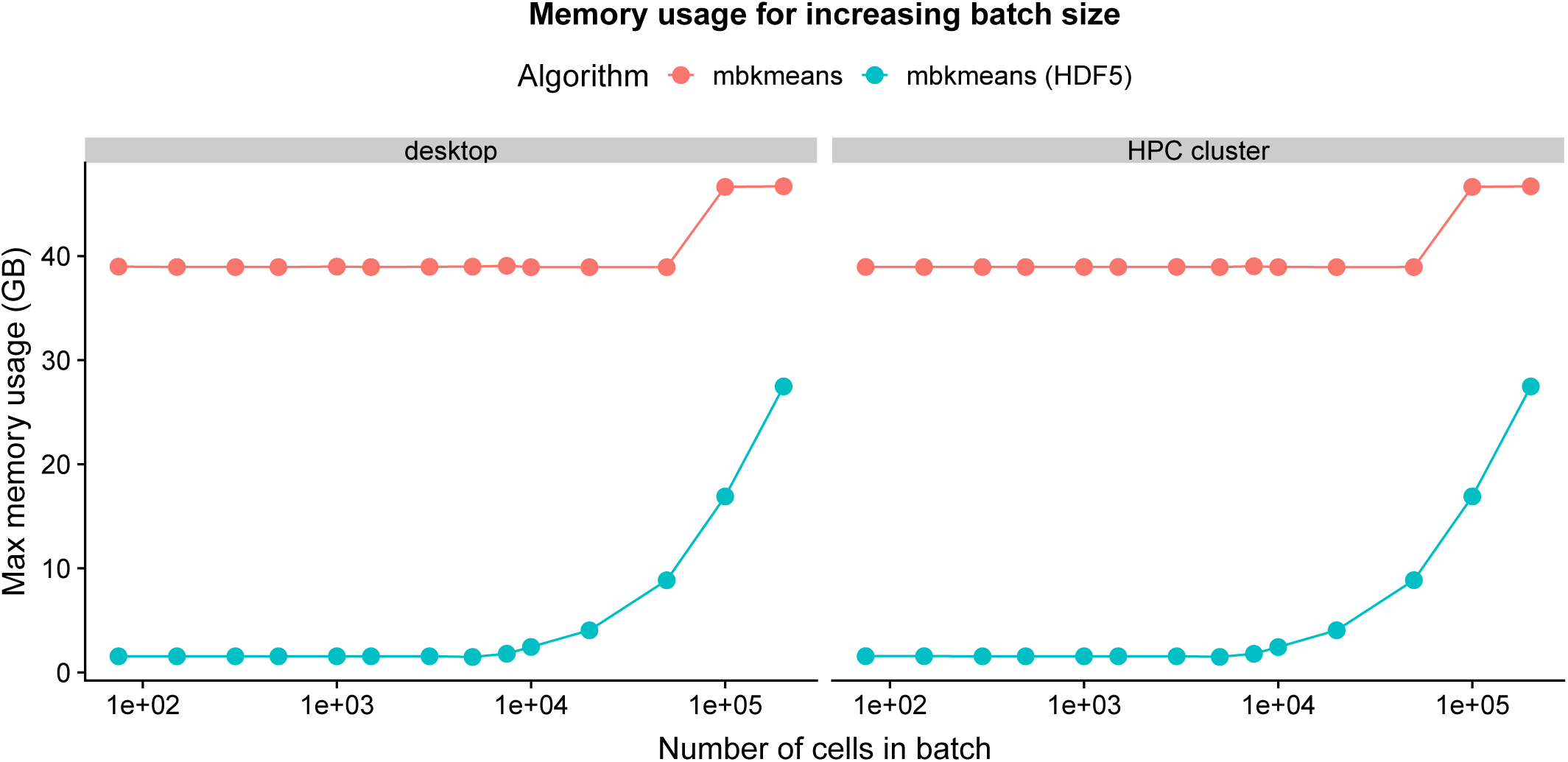
Memory-usage reported for both desktop and HPC cluster corresponding to Fig. 3.

**Fig. S11.**
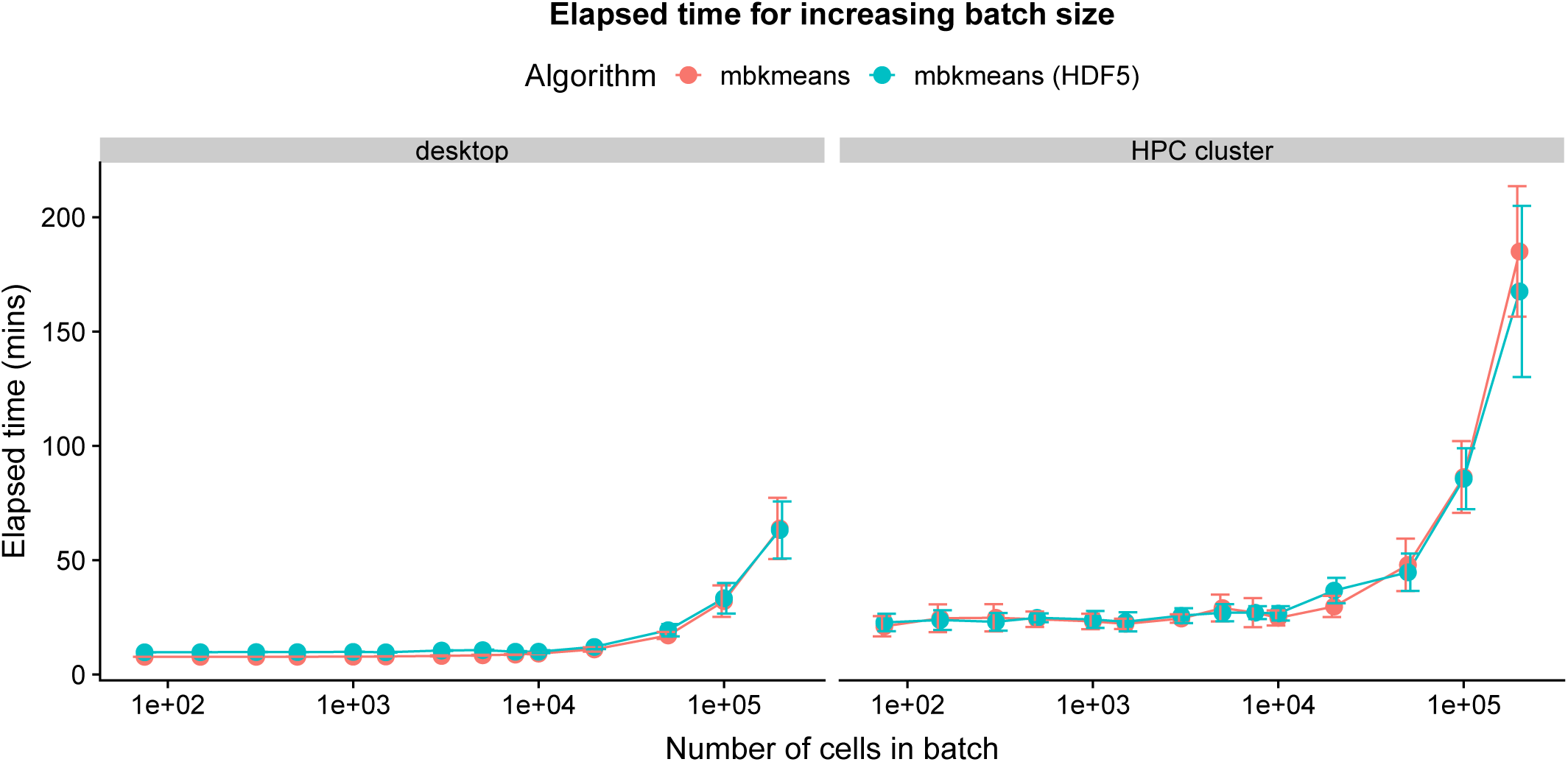
Elapsed time (minutes) reported for both desktop and HPC cluster corresponding to Fig. 3.

**Fig. S12.**
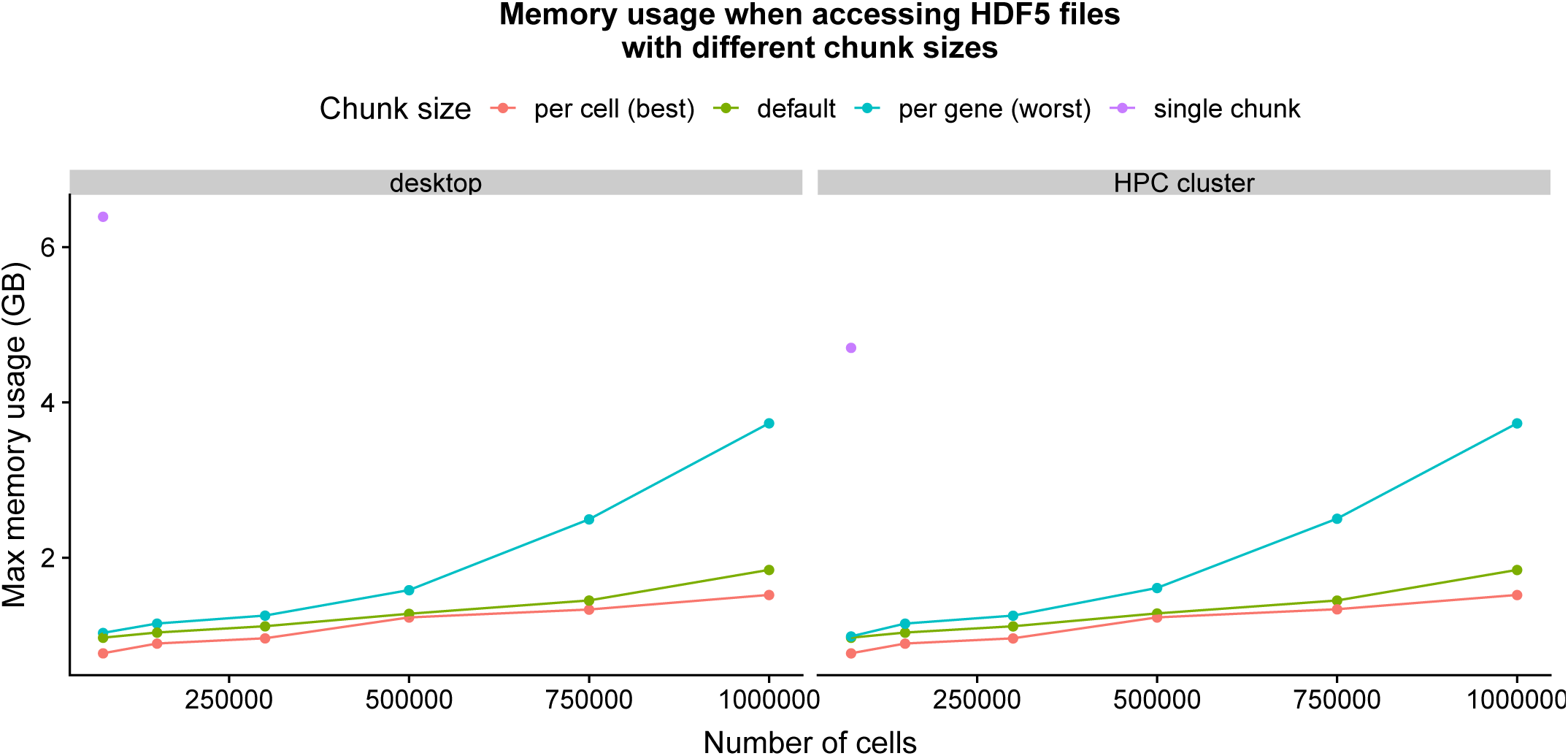
Memory-usage reported for both desktop and HPC cluster corresponding to Fig. 4.

**Fig. S13.**
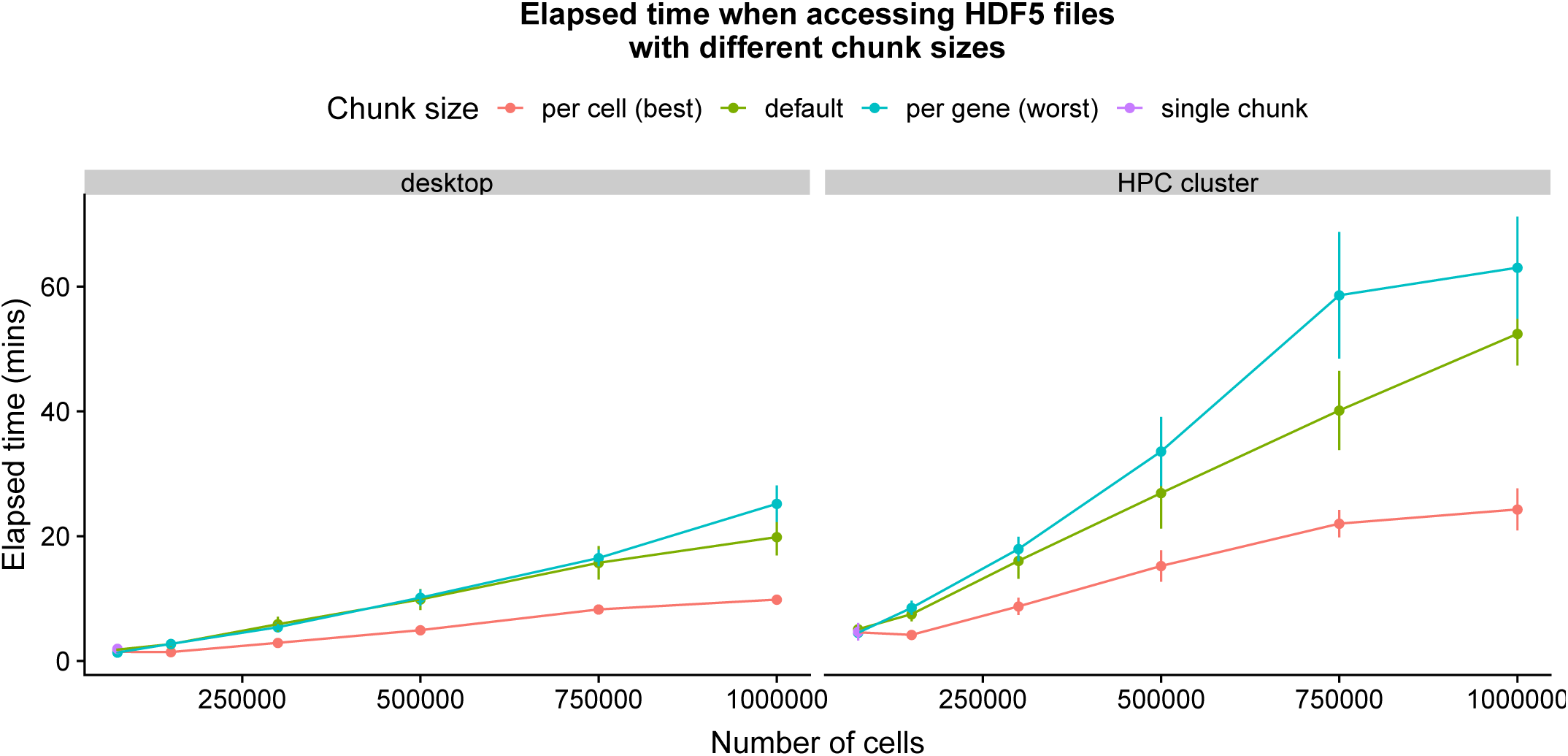
Elapsed time (minutes) reported for both desktop and HPC cluster corresponding to Fig. 4.

**Table S1.**
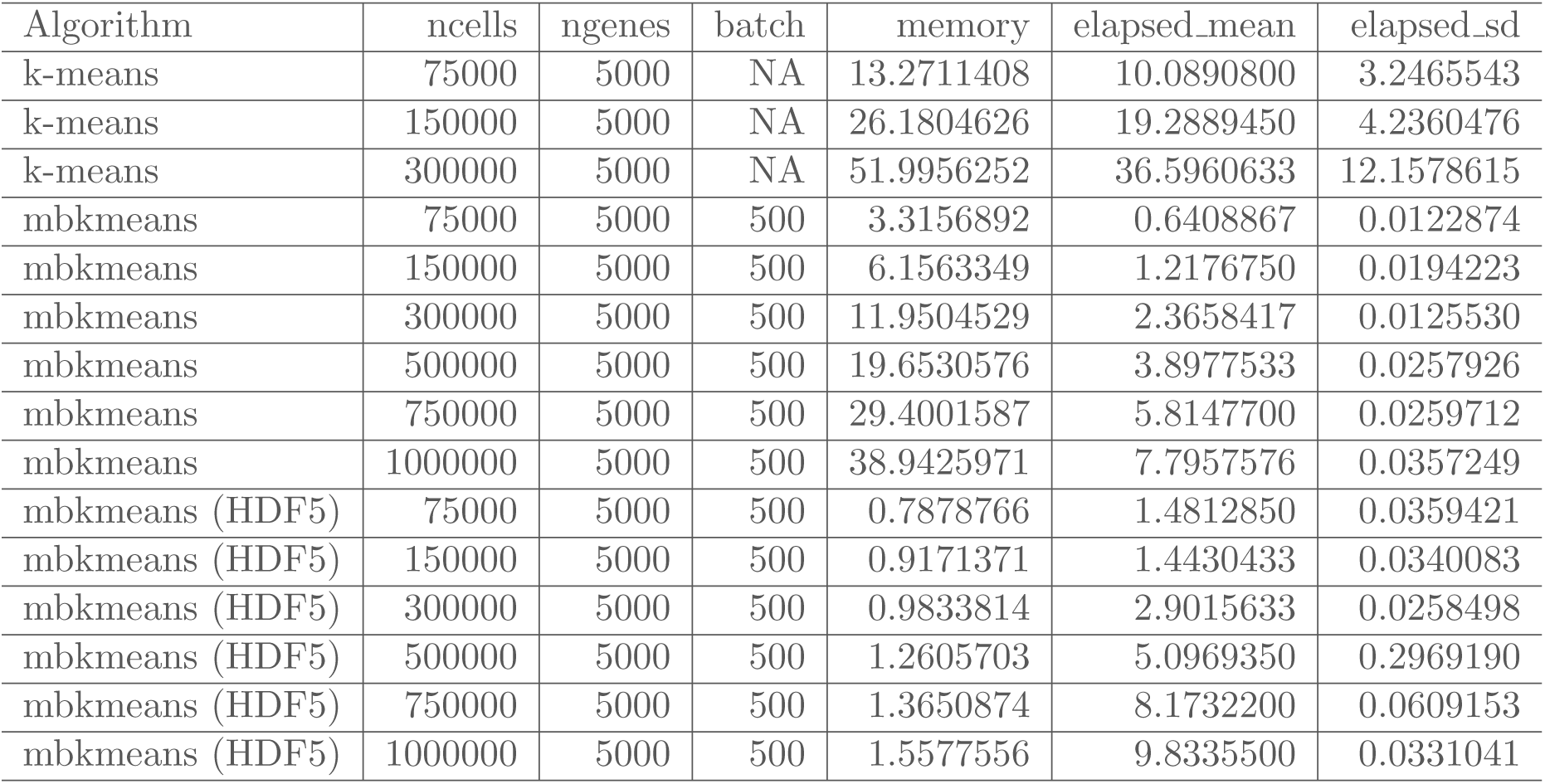
Performance evaluation for memory-usage and elapsed time as reported in Figure 1. We report the maximum memory (RAM) used (GB) and averaged elapsed time (minutes) for increasing sizes of datasets with *N* = 75,000, 150,000, 300,000, 500,000, 750,000, and 1,000,000 observations and 5,000 genes using a desktop computer. The average elapsed time (elapsed mean) and standard deviation (elapsed sd) of ten runs is reported in the table. We used *k* = 15 for both algorithms and used a batch size of *b* = 500 observations for *mbkmeans*.

| Algorithm | ncells | ngenes | batch | memory | elapsed_mean | elapsed_sd |
| --- | --- | --- | --- | --- | --- | --- |
| k-means | 75000 | 5000 | NA | 13.2711408 | 10.0890800 | 3.2465543 |
| k-means | 150000 | 5000 | NA | 26.1804626 | 19.2889450 | 4.2360476 |
| k-means | 300000 | 5000 | NA | 51.9956252 | 36.5960633 | 12.1578615 |
| mbkmeans | 75000 | 5000 | 500 | 3.3156892 | 0.6408867 | 0.0122874 |
| mbkmeans | 150000 | 5000 | 500 | 6.1563349 | 1.2176750 | 0.0194223 |
| mbkmeans | 300000 | 5000 | 500 | 11.9504529 | 2.3658417 | 0.0125530 |
| mbkmeans | 500000 | 5000 | 500 | 19.6530576 | 3.8977533 | 0.0257926 |
| mbkmeans | 750000 | 5000 | 500 | 29.4001587 | 5.8147700 | 0.0259712 |
| mbkmeans | 1000000 | 5000 | 500 | 38.9425971 | 7.7957576 | 0.0357249 |
| mbkmeans (HDF5) | 75000 | 5000 | 500 | 0.7878766 | 1.4812850 | 0.0359421 |
| mbkmeans (HDF5) | 150000 | 5000 | 500 | 0.9171371 | 1.4430433 | 0.0340083 |
| mbkmeans (HDF5) | 300000 | 5000 | 500 | 0.9833814 | 2.9015633 | 0.0258498 |
| mbkmeans (HDF5) | 500000 | 5000 | 500 | 1.2605703 | 5.0969350 | 0.2969190 |
| mbkmeans (HDF5) | 750000 | 5000 | 500 | 1.3650874 | 8.1732200 | 0.0609153 |
| mbkmeans (HDF5) | 1000000 | 5000 | 500 | 1.5577556 | 9.8335500 | 0.0331041 |

**Table S2.** Performance evaluation for accuracy as reported in Figure 2. We report the adjusted Rand index (ARI) and within-cluster sum of squares (WCSS) averaged across 50 replicates for increasing sizes of datasets with *N* = 5,000, 10,000, and 25,000 observations and increasing batch sizes *b* = 10, 35, 75, 150, 500, 750, 1,000 using a desktop computer. The average ARI for simulated data (ari sim mean) and standard deviation (ari sim sd), average WCSS for simulated data (wcss sim mean) and standard deviation (wcss sim sd), average WCSS for real scRNA-seq data (ari real mean) and standard deviation (ari real sd), is reported in the table. We used *k* = 3 for simulated data and *k* = 15 for real scRNA-seq data for all algorithms.

| Algorithm | ncells | batch | ari_sim_mean | ari_sim_sd | wcss_sim_mean | wcss_sim_sd | wcss_real_mean | wcss_real_sd |
| --- | --- | --- | --- | --- | --- | --- | --- | --- |
| k-means | 5000 | 10 | 0.9897752 | 0.0028632 | 1991.965 | 33.94950 | 23602.62 | 2763.306 |
| k-means | 5000 | 35 | 0.9894312 | 0.0023131 | 1994.014 | 43.30310 | 23603.06 | 2801.222 |
| k-means | 5000 | 75 | 0.9896092 | 0.0027030 | 1988.557 | 38.73366 | 23610.52 | 2963.918 |
| k-means | 5000 | 150 | 0.9891406 | 0.0023747 | 1986.485 | 32.85026 | 23613.23 | 2847.696 |
| k-means | 5000 | 500 | 0.9893209 | 0.0028165 | 1993.478 | 33.45500 | 23598.67 | 2859.045 |
| k-means | 5000 | 750 | 0.9886328 | 0.0027243 | 1986.465 | 30.49316 | 23597.09 | 2588.853 |
| k-means | 5000 | 1000 | 0.9893435 | 0.0026003 | 1991.071 | 26.37259 | 23597.66 | 2809.380 |
| k-means | 10000 | 10 | 0.9901098 | 0.0020059 | 1995.623 | 31.83414 | 23134.54 | 2675.893 |
| k-means | 10000 | 35 | 0.9891475 | 0.0021235 | 1988.504 | 25.67347 | 23134.54 | 2918.707 |
| k-means | 10000 | 75 | 0.9898653 | 0.0019818 | 1998.798 | 35.00856 | 23126.82 | 2718.442 |
| k-means | 10000 | 150 | 0.9896362 | 0.0018615 | 1989.612 | 28.83021 | 23072.22 | 2748.874 |
| k-means | 10000 | 500 | 0.9892754 | 0.0016966 | 1984.553 | 30.20538 | 23126.74 | 2757.786 |
| k-means | 10000 | 750 | 0.9893651 | 0.0019406 | 1991.718 | 27.53742 | 23072.42 | 2698.043 |
| k-means | 10000 | 1000 | 0.9888178 | 0.0019326 | 1992.960 | 27.05057 | 23130.62 | 2830.832 |
| k-means | 25000 | 10 | 0.9895777 | 0.0010857 | 1992.589 | 27.67066 | 26848.05 | 2457.034 |
| k-means | 25000 | 35 | 0.9896812 | 0.0013149 | 1988.434 | 28.06158 | 26847.19 | 2405.617 |
| k-means | 25000 | 75 | 0.9894319 | 0.0013192 | 1998.430 | 27.74529 | 26886.25 | 1915.763 |
| k-means | 25000 | 150 | 0.9896375 | 0.0011688 | 1988.218 | 29.89073 | 26802.16 | 2502.968 |
| k-means | 25000 | 500 | 0.9895120 | 0.0010132 | 1994.708 | 26.17106 | 26847.25 | 2400.455 |
| k-means | 25000 | 750 | 0.9895160 | 0.0011863 | 1994.466 | 25.20051 | 26886.50 | 2158.913 |
| k-means | 25000 | 1000 | 0.9893409 | 0.0010804 | 1989.884 | 27.91455 | 26848.05 | 2363.986 |
| mbkmeans | 5000 | 10 | 0.9791041 | 0.0149879 | 2787.158 | 268.53459 | 75926.52 | 36810.374 |
| mbkmeans | 5000 | 35 | 0.9874020 | 0.0046030 | 2190.102 | 56.82309 | 47257.80 | 13934.285 |
| mbkmeans | 5000 | 75 | 0.9881844 | 0.0043232 | 2076.647 | 52.05266 | 41642.46 | 8370.062 |
| mbkmeans | 5000 | 150 | 0.9897095 | 0.0030111 | 2031.929 | 38.99991 | 36584.62 | 6113.027 |
| mbkmeans | 5000 | 500 | 0.9897033 | 0.0034483 | 2005.876 | 34.42871 | 30958.60 | 3244.960 |
| mbkmeans | 5000 | 750 | 0.9892001 | 0.0025492 | 2002.357 | 35.44378 | 26249.06 | 3126.913 |
| mbkmeans | 5000 | 1000 | 0.9893587 | 0.0027131 | 1994.014 | 35.98165 | 26275.76 | 3167.850 |
| mbkmeans | 10000 | 10 | 0.9800142 | 0.0165777 | 2731.516 | 231.80159 | 66775.23 | 57543.709 |
| mbkmeans | 10000 | 35 | 0.9877258 | 0.0044289 | 2191.873 | 56.40113 | 54591.46 | 11713.384 |
| mbkmeans | 10000 | 75 | 0.9886893 | 0.0028865 | 2072.598 | 40.28502 | 38495.49 | 8133.570 |
| mbkmeans | 10000 | 150 | 0.9893979 | 0.0026309 | 2036.436 | 33.41415 | 34035.43 | 4781.714 |
| mbkmeans | 10000 | 500 | 0.9900648 | 0.0020835 | 2006.833 | 30.99967 | 30586.92 | 3621.350 |
| mbkmeans | 10000 | 750 | 0.9894255 | 0.0020135 | 2009.233 | 29.13326 | 26004.16 | 2621.284 |
| mbkmeans | 10000 | 1000 | 0.9895980 | 0.0020915 | 2005.894 | 37.90567 | 25230.06 | 2222.689 |
| mbkmeans | 25000 | 10 | 0.9819505 | 0.0102124 | 2681.955 | 217.64267 | 73934.80 | 37316.231 |
| mbkmeans | 25000 | 35 | 0.9877911 | 0.0038217 | 2184.486 | 68.01518 | 40138.51 | 8281.448 |
| mbkmeans | 25000 | 75 | 0.9886926 | 0.0024192 | 2075.565 | 34.98933 | 35590.41 | 5308.868 |
| mbkmeans | 25000 | 150 | 0.9895977 | 0.0020971 | 2048.228 | 29.86453 | 30315.25 | 3654.873 |
| mbkmeans | 25000 | 500 | 0.9894560 | 0.0013030 | 2005.893 | 27.30946 | 25393.64 | 2400.273 |
| mbkmeans | 25000 | 750 | 0.9896804 | 0.0012925 | 2003.314 | 28.53471 | 27233.51 | 2648.949 |
| mbkmeans | 25000 | 1000 | 0.9897113 | 0.0013482 | 1998.266 | 28.00118 | 25523.37 | 2613.946 |
| mbkmeans (HDF5) | 5000 | 10 | 0.9736589 | 0.0601490 | 2700.628 | 279.53992 | 92209.53 | 58576.452 |
| mbkmeans (HDF5) | 5000 | 35 | 0.9880798 | 0.0042995 | 2192.914 | 78.09871 | 50082.42 | 13021.588 |
| mbkmeans (HDF5) | 5000 | 75 | 0.9889910 | 0.0031843 | 2077.278 | 48.33555 | 40336.12 | 9929.519 |
| mbkmeans (HDF5) | 5000 | 150 | 0.9893773 | 0.0033271 | 2036.508 | 38.06441 | 35382.73 | 6623.128 |
| mbkmeans (HDF5) | 5000 | 500 | 0.9898477 | 0.0026405 | 2008.057 | 31.27833 | 28177.16 | 3662.996 |
| mbkmeans (HDF5) | 5000 | 750 | 0.9887941 | 0.0027125 | 2003.054 | 37.55634 | 26082.47 | 2652.004 |
| mbkmeans (HDF5) | 5000 | 1000 | 0.9889546 | 0.0026009 | 2005.522 | 32.72593 | 26161.00 | 3154.106 |
| mbkmeans (HDF5) | 10000 | 10 | 0.9815105 | 0.0109083 | 2677.676 | 267.91432 | 62060.41 | 36992.368 |
| mbkmeans (HDF5) | 10000 | 35 | 0.9882175 | 0.0045337 | 2202.861 | 85.99856 | 46748.12 | 14494.096 |
| mbkmeans (HDF5) | 10000 | 75 | 0.9877957 | 0.0033073 | 2089.668 | 46.65908 | 36736.58 | 6796.656 |
| mbkmeans (HDF5) | 10000 | 150 | 0.9895137 | 0.0022329 | 2029.635 | 28.50904 | 32702.95 | 4389.674 |
| mbkmeans (HDF5) | 10000 | 500 | 0.9893282 | 0.0023843 | 2000.400 | 30.03296 | 26901.48 | 2629.065 |
| mbkmeans (HDF5) | 10000 | 750 | 0.9894804 | 0.0020260 | 1998.409 | 21.70970 | 26291.87 | 2835.789 |
| mbkmeans (HDF5) | 10000 | 1000 | 0.9896635 | 0.0017741 | 1998.197 | 37.25141 | 25382.89 | 2874.408 |
| mbkmeans (HDF5) | 25000 | 10 | 0.9806960 | 0.0181099 | 2700.200 | 236.98421 | 62142.26 | 31252.643 |
| mbkmeans (HDF5) | 25000 | 35 | 0.9879255 | 0.0037378 | 2182.467 | 61.61046 | 40465.15 | 14752.192 |
| mbkmeans (HDF5) | 25000 | 75 | 0.9888413 | 0.0034479 | 2085.868 | 43.28525 | 36327.36 | 5801.333 |
| mbkmeans (HDF5) | 25000 | 150 | 0.9894029 | 0.0017897 | 2038.023 | 26.92411 | 31323.11 | 3406.756 |
| mbkmeans (HDF5) | 25000 | 500 | 0.9896004 | 0.0016466 | 2000.486 | 19.19524 | 27258.32 | 2672.190 |
| mbkmeans (HDF5) | 25000 | 750 | 0.9897985 | 0.0011803 | 1993.182 | 26.25207 | 25686.95 | 2587.394 |
| mbkmeans (HDF5) | 25000 | 1000 | 0.9895056 | 0.0011316 | 1996.803 | 32.75242 | 26705.52 | 2546.067 |

**Table S3.**
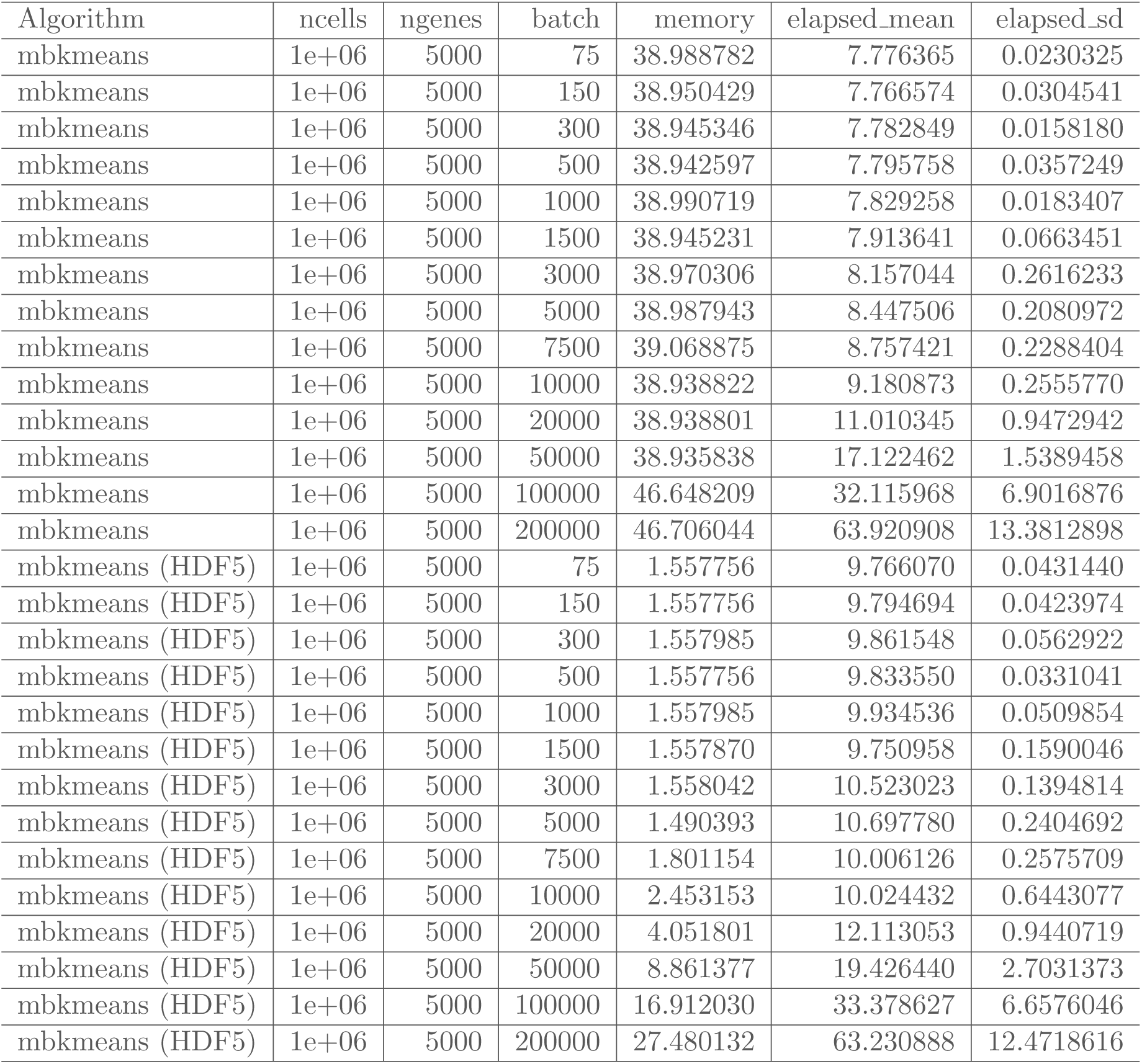
Performance evaluation for memory-usage and elapsed time reported in Figure 3. We report the maximum memory (RAM) used (GB) and averaged elapsed time (minutes) for increasing batch sizes with *b* = 75, 150, 300, 500, 1,000, 1,500, 3,000, 5,000, 7,500, 10,000, 20,000, 50,000, 100,000, 200,000 with a dataset of size *N* = 1,000,000 observations and 5,000 genes using a desktop computer. The average (elapsed mean) and standard deviation (elapsed sd) of ten runs is reported in the table. We used *k* = 15 for the number of centroids in *mbkmeans*.

| Algorithm | ncells | ngenes | batch | memory | elapsed_mean | elapsed_sd |
| --- | --- | --- | --- | --- | --- | --- |
| mbkmeans | 1e+06 | 5000 | 75 | 38.988782 | 7.776365 | 0.0230325 |
| mbkmeans | 1e+06 | 5000 | 150 | 38.950429 | 7.766574 | 0.0304541 |
| mbkmeans | 1e+06 | 5000 | 300 | 38.945346 | 7.782849 | 0.0158180 |
| mbkmeans | 1e+06 | 5000 | 500 | 38.942597 | 7.795758 | 0.0357249 |
| mbkmeans | 1e+06 | 5000 | 1000 | 38.990719 | 7.829258 | 0.0183407 |
| mbkmeans | 1e+06 | 5000 | 1500 | 38.945231 | 7.913641 | 0.0663451 |
| mbkmeans | 1e+06 | 5000 | 3000 | 38.970306 | 8.157044 | 0.2616233 |
| mbkmeans | 1e+06 | 5000 | 5000 | 38.987943 | 8.447506 | 0.2080972 |
| mbkmeans | 1e+06 | 5000 | 7500 | 39.068875 | 8.757421 | 0.2288404 |
| mbkmeans | 1e+06 | 5000 | 10000 | 38.938822 | 9.180873 | 0.2555770 |
| mbkmeans | 1e+06 | 5000 | 20000 | 38.938801 | 11.010345 | 0.9472942 |
| mbkmeans | 1e+06 | 5000 | 50000 | 38.935838 | 17.122462 | 1.5389458 |
| mbkmeans | 1e+06 | 5000 | 100000 | 46.648209 | 32.115968 | 6.9016876 |
| mbkmeans | 1e+06 | 5000 | 200000 | 46.706044 | 63.920908 | 13.3812898 |
| mbkmeans (HDF5) | 1e+06 | 5000 | 75 | 1.557756 | 9.766070 | 0.0431440 |
| mbkmeans (HDF5) | 1e+06 | 5000 | 150 | 1.557756 | 9.794694 | 0.0423974 |
| mbkmeans (HDF5) | 1e+06 | 5000 | 300 | 1.557985 | 9.861548 | 0.0562922 |
| mbkmeans (HDF5) | 1e+06 | 5000 | 500 | 1.557756 | 9.833550 | 0.0331041 |
| mbkmeans (HDF5) | 1e+06 | 5000 | 1000 | 1.557985 | 9.934536 | 0.0509854 |
| mbkmeans (HDF5) | 1e+06 | 5000 | 1500 | 1.557870 | 9.750958 | 0.1590046 |
| mbkmeans (HDF5) | 1e+06 | 5000 | 3000 | 1.558042 | 10.523023 | 0.1394814 |
| mbkmeans (HDF5) | 1e+06 | 5000 | 5000 | 1.490393 | 10.697780 | 0.2404692 |
| mbkmeans (HDF5) | 1e+06 | 5000 | 7500 | 1.801154 | 10.006126 | 0.2575709 |
| mbkmeans (HDF5) | 1e+06 | 5000 | 10000 | 2.453153 | 10.024432 | 0.6443077 |
| mbkmeans (HDF5) | 1e+06 | 5000 | 20000 | 4.051801 | 12.113053 | 0.9440719 |
| mbkmeans (HDF5) | 1e+06 | 5000 | 50000 | 8.861377 | 19.426440 | 2.7031373 |
| mbkmeans (HDF5) | 1e+06 | 5000 | 100000 | 16.912030 | 33.378627 | 6.6576046 |
| mbkmeans (HDF5) | 1e+06 | 5000 | 200000 | 27.480132 | 63.230888 | 12.4718616 |

**Table S4.** Performance evaluation for memory-usage and elapsed time as reported in Figure 4. We report the maximum memory (RAM) used (GB) and averaged elapsed time (minutes) for increasing sizes of datasets with *N* = 75,000, 150,000, 300,000, 500,000, 750,000, and 1,000,000 observations and 5,000 genes using a desktop computer. The average elapsed time (elapsed mean) and standard deviation (elapsed sd) of ten runs is reported in the table. The single chunk was only able to run for the smallest dataset size (*N* = 75,000). We used *k* = 15 and used a batch size of *b* = 500 observations.

| chunk_size | dim_1 | dim_2 | ncells | ngenes | batch | memory | elapsed_mean | elapsed_sd |
| --- | --- | --- | --- | --- | --- | --- | --- | --- |
| per cell (best) | 5000 | 1 | 75000 | 5000 | 500 | 0.7693644 | 1.463833 | 0.0335315 |
| per cell (best) | 5000 | 1 | 150000 | 5000 | 500 | 0.8950103 | 1.434578 | 0.0219519 |
| per cell (best) | 5000 | 1 | 300000 | 5000 | 500 | 0.9622431 | 2.909230 | 0.0384201 |
| per cell (best) | 5000 | 1 | 500000 | 5000 | 500 | 1.2307574 | 4.941737 | 0.0175054 |
| per cell (best) | 5000 | 1 | 750000 | 5000 | 500 | 1.3326299 | 8.265250 | 0.2912552 |
| per cell (best) | 5000 | 1 | 1000000 | 5000 | 500 | 1.5210066 | 9.840213 | 0.3331764 |
| default | 258 | 3872 | 75000 | 5000 | 500 | 0.9702795 | 1.822533 | 0.1901076 |
| default | 182 | 5477 | 150000 | 5000 | 500 | 1.0366168 | 2.709267 | 0.4417234 |
| default | 129 | 7745 | 300000 | 5000 | 500 | 1.1176969 | 5.901668 | 1.1833504 |
| default | 100 | 10000 | 500000 | 5000 | 500 | 1.2780040 | 9.865945 | 1.6815279 |
| default | 81 | 12247 | 750000 | 5000 | 500 | 1.4494440 | 15.730765 | 2.6753175 |
| default | 70 | 14142 | 1000000 | 5000 | 500 | 1.8435055 | 19.860840 | 2.9493403 |
| per gene (worst) | 1 | 75000 | 75000 | 5000 | 500 | 1.0318118 | 1.359693 | 0.1138983 |
| per gene (worst) | 1 | 150000 | 150000 | 5000 | 500 | 1.1522158 | 2.741733 | 0.3571907 |
| per gene (worst) | 1 | 300000 | 300000 | 5000 | 500 | 1.2548324 | 5.412373 | 0.4938202 |
| per gene (worst) | 1 | 500000 | 500000 | 5000 | 500 | 1.5828893 | 10.148773 | 1.3119979 |
| per gene (worst) | 1 | 750000 | 750000 | 5000 | 500 | 2.4946825 | 16.504955 | 1.3132680 |
| per gene (worst) | 1 | 1000000 | 1000000 | 5000 | 500 | 3.7318117 | 25.202253 | 2.9298333 |
| single chunk | 5000 | 75000 | 75000 | 5000 | 500 | 6.3917671 | 1.967600 | NA |

**Table S5.**
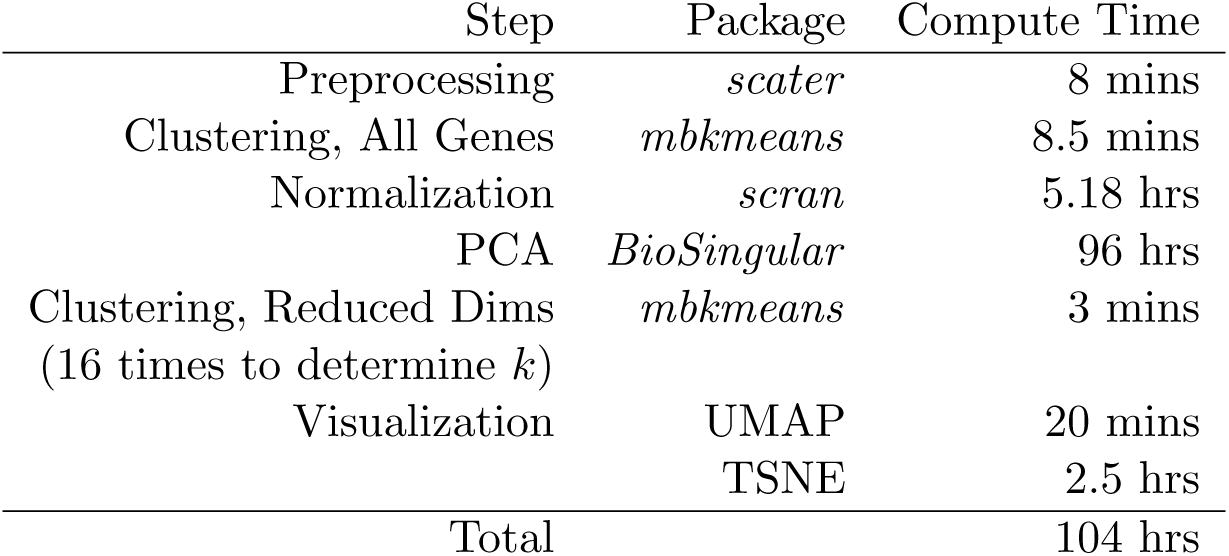
Computational time for each of the steps of the pipeline for the full 1.3 million mouse brain cells

**Table S6.** Identification of *mbkmeans* clusters with Marker Genes

| *mbkmeans* Cluster | Genes | Biological Subtype | References |
| --- | --- | --- | --- |
| 1 | *Mdk, Mki67, Nde1* | Radial precursor / Proliferating | [44, 41] |
| 2 | *Gad1, Gad2, Foxp2, Syt6, Bcl11b* | Interneurons | [45, 43, 41] |
| 3 | *Neurod1, Synpr, Prox1* | Granule Cells | [40] |
| 4 | *Sox4, Sox11, Meis2* | Pyramidal progenitors | [40] |
| 5 | *Gad1, Gad2, Sst, Lhx6* | Interneurons | [45, 43, 41] |
| 6 | *Meis2, Neurod6* | Pyramidal progenitors | [40] |
| 7 | *Vim, Fabp7, Sox9, Dbi* | Radial precursor | [44] |
| 8 | *Crym, Tnr, Cck* | Pyramidal neurons | [41, 45] |
| 9 | *Sstr2, Mdk, Eomes* | Neuronal progenitors | [44, 41] |
| 10 | *Pdgfra, Vim, Dbi* | OPC | [40] |
| 11 | *Cck, Ntf3, Nr4a2* | Pyramidal neurons | [40, 43, 45] |
| 12 | *Rgs5, Igfbp7* | Endothelial/SMC/Mural | [40, 41, 42] |
| 13 | *Neurod6, Satb2, Meis2* | Pyramidal progenitors | [40] |
| 14 | *Hbbbs, Hbaa1, Hbaa2* | Blood |  |
| 15 | *Reln, Pcp4, Tbr1* | Cajal-Retzius cells | [46] |

